# Something old, something new: the origins of an unusual renal cell underpinning a beetle water-conserving mechanism

**DOI:** 10.1101/2024.03.01.582930

**Authors:** Robin Beaven, Takashi Koyama, Muhammad T. Naseem, Kenneth V. Halberg, Barry Denholm

## Abstract

Tenebrionid beetles have been highly successful in colonising environments where water is scarce, underpinned by their unique osmoregulatory adaptations. These include a cryptonephridial arrangement of their organs, in which part of their renal/Malpighian tubules are bound to the surface of the rectum. This allows them to generate a steep osmotic gradient to draw water from within the rectum and return it to the body. Within the cryptonephridial tubules a seemingly novel cell type, the leptophragmata, is considered to play a key role in transporting potassium chloride to generate this osmotic gradient. Nothing was known about the developmental mechanisms or evolution of these unusual renal cells. Here we investigate the mechanisms underpinning development of the leptophragmata in the red flour beetle, *Tribolium castaneum*. We find that leptophragmata express and require a *teashirt*/*tiptop* transcription factor gene, as do the secondary renal cells of *Drosophila melanogaster* which lack a cryptonephridial arrangement. We also find an additional transcription factor, Dachshund, is required to establish leptophragmata identity and to distinguish them from the secondary cells in *Tribolium’s* non-cryptonephridial region of renal tubule. Dachshund is also expressed in a sub-population of secondary cells in *Drosophila*. So leptophragmata, which are unique to the beetle lineage, appear to have originated from a specific renal cell type present ancestrally, and specified by a conserved repertoire of transcription factors.

**Significance:** Beetles are a highly successful insect group and represent a quarter of all known animal species. Their digestive/renal systems have undergone major evolutionary change compared to other insects, likely contributing to their success. A dramatic example is the cryptonephridial complex, an evolutionary innovation of the gut and renal system which integrate as a powerful water-conservation system; an adaptation for survival in arid conditions. An unusual renal cell type—the leptophragmata—underpin the functions of the complex, but their developmental and evolutionary origins are unknown. Here we reveal the developmental mechanism that establish leptophragmata identity and, by studying a species lacking a cryptonephridial complex, shed light on their evolutionary origin. More broadly, the work illuminates the evolution of novel cell types.

## Introduction

Tenebrionid beetles have been extremely successful at surviving where water is scarce, and therefore colonising arid regions (1). Unique osmoregulatory adaptations are likely to underpin this ability, including differences in the physiology and hormonal control of their renal/Malpighian tubules (MpTs) compared to other insects (2–4). Another critical adaptation is the cryptonephridial complex (CNC). In contrast to the ancestral state, where the MpTs are free within the haemolymph, in the CNC arrangement the distal portions of the MpTs are bound to the surface of the rectum (5). This complex is ensheathed in an insulating tissue, the perinephric membrane (6). Ion transport by the MpTs generates a high potassium chloride concentration surrounding the rectum (7), which functions to draw water osmotically out from the faeces within the rectal lumen, which can then be returned by the MpTs to the body (8). This allows tenebrionid beetles such as *Tenebrio molitor* to generate powder dry faeces, so minimising water loss (6, 9). The ability of the CNC to extract water from the rectum is so powerful that they can even use it to extract moisture from the air in conditions above a certain humidity (10–13).

Leptophragmata are a seemingly novel cell type within the MpTs of the CNC, in terms of their specialised position, morphology and physiological function, and are central to how the CNC works. They have unique access to the haemolymph surrounding the CNC, residing beneath regions of the perinephric membrane where it is present as only an extremely thin ‘blister’ of extracellular material (6, 14). Leptophragmata have long been considered the likely route by which chloride ions enter the CNC (6, 14, 15), although other possibile routes have been suggested, including from the fluid in the rectal lumen (7). Using the model insect species the red flour beetle (*Tribolium castaneum*) we have shed light on the molecular mechanisms underpinning leptophragmata function. We identified a cation/proton antiporter, NHA1, in the leptophragmata, which likely uses a proton gradient to transport potassium into the CNC (16). In turn, chloride is considered to follow along the electrochemical gradient established by potassium transport (7, 8).

We previously demonstrated that *Tribolium* has a CNC structure very similar to that of *T. molitor*, in which the CNC has been most studied. *Tribolium* is an ideal species to study CNC developmental mechanisms and their molecular basis (16, 17), having a sequenced, well annotated genome, and established means for manipulating gene expression including RNAi. We previously identified Tiptop, a transcription factor of the *teashirt* gene family, to be expressed in *Tribolium* leptophragmata (17). Teashirt and its paralogue Tiptop are required for differentiation of MpT secondary cells in *Drosophila* (18). *Tribolium* Tiptop is expressed in leptophragmata but also in the free tubule (the region of the tubule not associated with the rectum and exposed to the haemolymph) in a subset of cells which display properties akin to *Drosophila* secondary cells, including smaller nuclear size (17). Tiptop is the sole member of the Teashirt/Tiptop family of transcription factors in *Tribolium* (19).

Here we build on these insights to gain an understanding of the molecular mechanisms by which leptophragmata identity is specified, and provide clues of their evolutionary origins. We draw on the tissue specific transcriptomic database, BeetleAtlas, which has proven a powerful resource for guiding molecular studies in *Tribolium* (16, 20), and conduct functional studies using the genetic manipulation, cellular morphological and physiological methods which are established for this species. We find that the transcription factor *tiptop*, in combination with *dachshund*, are required for leptophragmata identity. We also find evidence that this combination of genes defines a specific sub-population of secondary cells in *Drosophila*, which lack a CNC. Therefore although differentiated leptophragmata appear unique to the beetle lineage at a morphological and molecular level, they are specified by a conserved repertoire of transcription factors, and therefore likely derived from a specialised renal cell type present ancestrally. This work sheds light on the evolution of a novel cell type within the CNC which underpins the remarkable water conservation ability of tenebrionid beetles.

## Methods

### Insect culturing and sample preparation

*Drosophila melanogaster* were cultured on standard media at 25°C*. w^1118^ Drosophila* embryos were collected on grape juice agar plates with yeast paste. Embryos were dechorionated in 50% bleach for ∼4 minutes, rinsed in water and fixed for ∼25 minutes in 4% formaldehyde in PBS / heptane on a rotor. 4% formaldehyde was replaced by methanol and vitelline membranes removed by shaking, before rinsing embryos in methanol. Embryos were then processed or stored in methanol at −20°C. Adult MpTs were dissected in PBS and fixed for ∼20 minutes in 4% formaldehyde in PBS. The following stocks were used: *tsh-c724-Gal4* (21), *CD8-GFP* (22), *UAS-dac* (3^rd^) (23), *UAS-dac-shRNA* (dac^HMS01435^, Bloomington # 35022), and *w*; UAS-GFP(Cytosolic); Clc-A-GAL4* (from Julian Dow and Anthony Dornan. *Clc-A-GAL4* is VDRC_ID 202625).

The *Tribolium castaneum* San Bernardino (SB) strain was used, with cultures maintained on wholemeal flour with yeast. Embryos were collected by allowing adults to lay on white flour for 3 days at 30°C, before removing embryos with a 300μm gauge sieve. Embryos were fixed as detailed for *Drosophila* embryos (above) with the following modification for removal of vitelline membranes: embryos were briefly rinsed in heptane and transferred onto double sided tape with a paintbrush. They were then bathed in PBS and removed from their vitelline membranes by hand using a pulled glass needle.

For larval/adult preparations of the CNC, guts with associated MpTs were dissected in PBS by pulling the posterior tip of the larval/adult abdomen away from the body which exposes the gut/CNC/MpTs, before fixing for ∼20 minutes in 4% formaldehyde in PBS. In some cases, in order to avoid damaging attachments between the MpTs and the gut, dissection was performed more carefully by first opening up the abdomen of the adult by cutting the cuticle with spring scissors (Fine Science Tools, 15000-08).

### dsRNA knockdown

For knockdown, dsRNA targeting specific genes was generated as follows. In most cases templates were amplified from genomic DNA. This was extracted from 5 adult beetles by freezing in 200μl lysis buffer (100mM Tris-HCl pH9, 100mM EGTA, 1% SDS) for 30 minutes before thawing and homogenising. A further 600μl of lysis buffer was added and the sample was incubated at 70°C for 30 minutes. 150μl of cooled 8M potassium acetate was added and incubated on ice for 20 minutes. The sample was then centrifuged (20 minutes, 13,000 rpm, 4°C). The equivalent of 0.9 of the supernatant volume, of cold isopropanol, was added to the supernatant to precipitate the DNA and centrifuged for 10 minutes (13,000 rpm, 4°C), the pellet was washed with 70% ethanol and air dried. The pellet was resuspended in 100μl TE buffer (QIAGEN) and 100μl of Phenol-chloroform was added, vortexed, and centrifuged for 15 minutes (13,000 rpm, 4°C). The upper phase was removed, and to this 5μl of 3M sodium acetate and 250μl of cold 100% ethanol were added, and left at −20°C overnight. The sample was centrifuged for 15 minutes (13,000 rpm, 4°C), the pellet washed in 70% ethanol and air dried. The pellet was finally resuspended in TE buffer. For *GFP* and *Ampicillin* (*Amp*) dsRNA, plasmids containing the relevant sequence were used to generate the template. Templates were generated by PCR, using primers to which *T7* promoter sequences had been added at the 5’ ends. The following primers were used (Table 1). dsRNA was synthesised from these templates using the MEGAscript T7 transcription kit (Invitrogen) and precipitated using LiCl. dsRNA pellets were finally resuspended in injection buffer (1.4mM NaCl, 0.07mM Na_2_HPO_4_, 0.03mM KH_2_PO_4_, 4mM KCl), and the approximate concentration of dsRNA was determined using a nano-drop, measuring optical density using a factor of 45 ng-cm/μl. The following concentrations of dsRNA were injected: *tio* (1) - 200ng/ μl, *tio* (2) - 850-3700ng/μl, *Pph13* - 1100ng/μl, *tup* - 1200-1800ng/μl, *eve* - 1975ng/μl, *twi* – 6151ng/μl, *elB* - 970ng/μl, *net* - 445ng/μl, *en-like* - 1480ng/μl, *SoxD* - 4195ng/μl, *disco* - 3750ng/μl, *dac* - 440-4440ng/μl, *GFP* - 3500ng/μl, *Amp* - 1350ng/μl. For *tio* dsRNA experiments either *tio* (2) or a mixture of *tio* (1) and *tio* (2) were used, which are previously used target sequences (19).

**Table 1.**
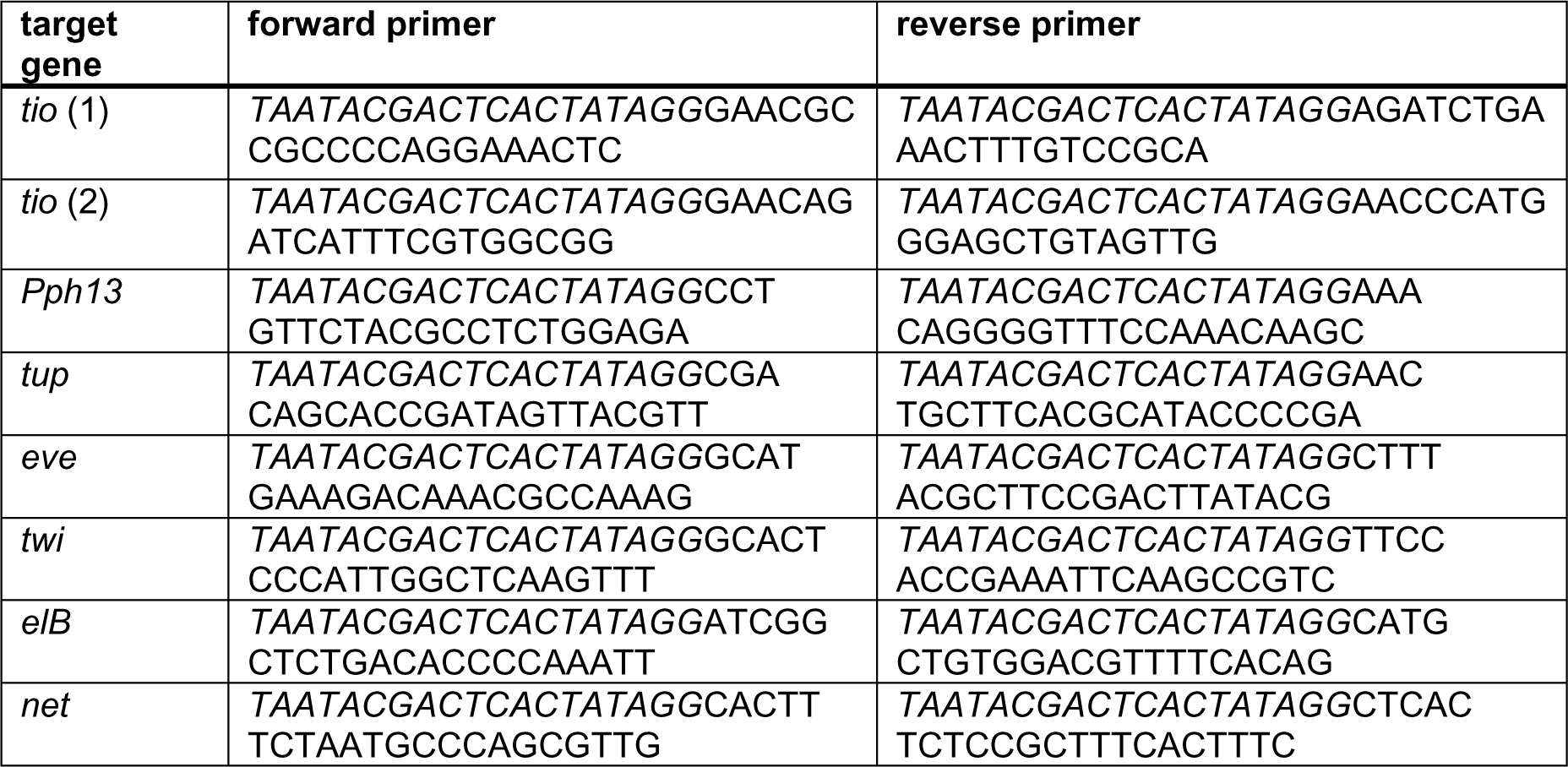
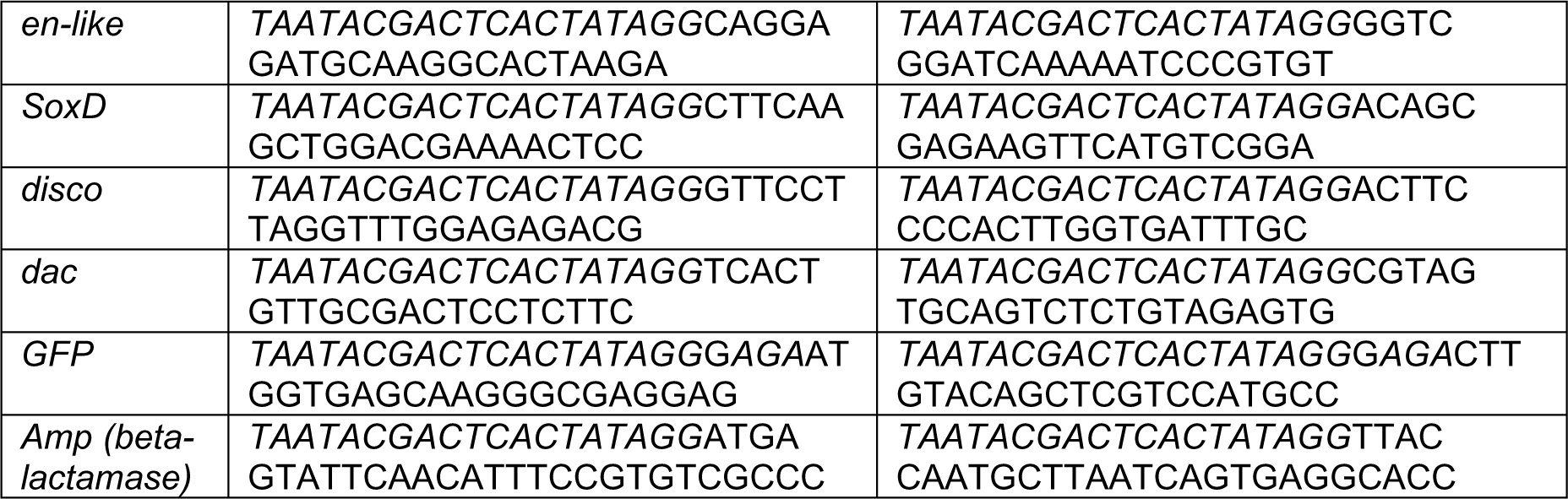
primer sequences for generation of templates for dsRNA synthesis. T7 sequence indicated in italic.

For maternal injection, female beetles were glued on their backs with Marabu Fixogum rubber cement. Pulled glass capillaries (Narishige GD-1, 1×90mm) with the tips broken were mounted in a needle manipulator and dsRNA was front-loaded using a syringe. Beetles were anaesthetised using CO_2_ and injected close to the posterior end in the softer cuticle beneath the elytra. Injected females together with uninjected males were placed on white flour with yeast. After a few days they were moved to fresh flour and eggs laid were collected and allowed to hatch into larvae. For larval injections, final instar larvae were injected as for adult females, except they were glued down ventrally and injected dorsally into the softer cuticle between body segments. Injected larvae were placed on sieved wholemeal flour with yeast and allowed to mature into adults.

### Immunohistochemistry and *in situ* hybridisation

For immunohistochemistry PBST + BSA (PBS with 0.3% Triton X-100 and 0.5% bovine serum albumin) was used to dilute antibodies and perform wash steps. The following antibodies were used: anti-Urn8R (rabbit, 1:200) (4), anti-Tio (rat, 1:100) (24), anti-Tsh (rabbit, 1:3000, from S. Cohen, IMCB, Singapore), anti-Dac (mouse, 1:100, mAbdac1-1-c from DSHB), anti-alpha-Tubulin (mouse, 1:20, AA4.3, DSHB), anti-GFP (goat, 1:500, ab6673, Abcam), anti-Clc-a (rabbit, 1:2000) (25), and anti-NHA1 (polyclonal rabbit, 1:500) (16). Secondary antibodies from Jackson ImmunoResearch of appropriate species tagged with Alexa Fluor 488, Cy3 or Cy5 fluorophores were used at 1:200, DAPI (Molecular Probes) was used 1:1000, and Alexa Fluor 488-568- or 647-conjugated phalloidin (Life Technologies) were used 1:200. Samples were mounted in 85% glycerol, 2.5% propyl gallate.

Fluorescent *in situ* hybridisation was carried out using the hybridisation chain reaction v3.0 (HCR v3.0) technique (26), with a probe set size of 20 for *cut*, *tio* and *dac* (Molecular Instruments). A protocol from Bruce and colleagues was used (27). Samples were mounted in 85% glycerol, 2.5% propyl gallate.

### Sliver staining assay

For the silver staining assay, guts were dissected in water and transferred to 1% silver nitrate in water, where they were incubated in the light for ∼4 minutes. They were then rinsed in water before fixing for ∼20 minutes in 4% formaldehyde in PBS. Samples were rinsed in PBS and mounted in DPX new (Merck).

### Microscopy and image analysis

Fluorescence images were taken using either a Nikon A1R or Zeiss LSM800 confocal microscope. Maximum intensity projection images were generated using Fiji. Samples from the silver staining were imaged with a Zeiss Axioplan light microscope with a Leica DFC 425C camera. To get an indication of the size of leptophragmata nuclei, the area was measured from a single z-section at approximately the midpoint of the nucleus. The perimeter of the nucleus was defined manually, and the area calculated using the freehand selections tool in Fiji. To calculate the staining intensity of *dac* mRNA from HCR samples images were take using equivalent settings. The region of the nucleus was defined in a single z-section, using the *tio* stained channel, and the mean staining intensity in the *dac* channel was calculated using Fiji.

### Ligand-Receptor Binding Assay

A synthetic analogue of Diuretic Hormone 37 (DH37) with an N-terminal cysteine was synthesized by Cambridge Peptides (Birmingham, UK) at a purity of >90%, in order to conjugate a TMR-C_5_-maleimide Bodipy dye (BioRad, CA, USA), to make fluorescent TMR-C_5_-maleimide-SPTISITAPIDVLRKTWAKENMRKQMQINREYLKNLQamide (DH37-F). The ex vivo receptor-binding assay was performed as described in (3, 4). Perirectal tubules attached to the perinephric membrane, were carefully dissected from the rectum under Schneider’s and *Tribolium* saline (1:1) and then mounted on poly-L-lysine-covered 35mm glass bottom dishes. Next, the tissues were set-up in a matched-pair protocol, in which one batch was incubated in the appropriate insect saline added the labelled neuropeptide analogue (10^-6^ M) and DAPI (1 µg/ml), while the other was incubated in just DAPI; the latter batch was used to adjust baseline filter and exposure settings to minimize auto-fluorescence during image acquisition. Images were subsequently recorded on a Zeiss LSM 900 confocal microscope using baseline filter and exposure settings.

### Physiological assays

For the physiological assays either 1 µg/µl *dac* dsRNA or *Amp* dsRNA (control) were injected into age-matched final larval instar larvae using a Nanoject II injector (Drummond Scientific, PA, USA). Animals 10-14 days after eclosion were used for experiments. To assess tolerance to desiccation, animals were transferred to a 96-well plate in a container filled with silica gel beads (Sigma-Aldrich, MO, USA) to produce a low-humidity environment (approx. R.H. 5%, measured by a custom-build hygrometer). Mortality rates of the treated adults were assessed over a 16-day period by viewing under a stereo microscope (4).

To assess the effects of Dachshund knockdown on excretory behaviour in intact living animals, dsRNA-injected animals were starved for 2 days followed by re-feeding a standard *Tribolium* medium supplemented with 0.05% (w/w) Bromophenol blue (BPB) sodium salt (Sigma-Aldrich, MO, USA) overnight as in (4, 16). This was created by mixing the standard *Tribolium* medium with BPB and a small amount of water to create a uniform paste, which was left to dry at room temperature overnight. The dried BPB-labelled *Tribolium* medium was then ground to a fine powder creating a consistency similar to that of the standard medium. Beetles were then placed into individual wells of a 96-well plate fitted with a small piece of filter paper and the number of BPB-labelled deposits produced by each animal over a 4 hour period was quantified. Next, the treated adults were recovered and raised in the same BPB medium for an additional 7 days, after which the number animals exhibiting defective defecation (i.e. whose anus was blocked by excretory product likely due to excess water content) was quantified under a stereo microscope.

For the *ex-vivo* fluid reabsorption assay, water reabsorption rates from the rectal complex were measured using a protocol described in (16). The control and *dachshund* knockdown animals were dissected carefully, keeping the head, gut, tubules and CNC intact under Schneider’s medium. MpTs were then broken from the entry point (common trunk) to the CNC. The dissected animals were carefully transferred to another dish containing paraffin oil (molecular grade) with a wax layer at the bottom, and gently stretched (to keep intact the fore-, mid-, hind-gut, MpTs, and CNC) and pinned at the head and anal cuticle. Two drops (5μl and 0.3μl) of premixed 3x *Tribolium* saline + Schneider’s solution (1:1) supplemented with 100 µmol/l of amaranth (Sigma-Aldrich, St Louis, MO, USA) were dropped under paraffin oil and their diameter measured. The 5μl drop was coaxed into place around the midgut and MpTs and the 0.3μl drop around the rectal complex with the help of a capillary pulled glass rod, ensuring contact with the head and anal cuticle was avoided. MpTs were gently drawn out from the saline drop and then the open ends of MpTs were wrapped around the pin with the help of fine forceps. The preparation was left for two hours—the maximum time the system remained stable. The change in volume of the initial and final drop was measured as volume (v) = (π x diameter^3^)/6, and the rate of water absorbance by the CNC was calculated by the following formula: J_fluid_ =Δv/Δt where J_fluid_ is the fluid reabsorption rate (nl/min), Δv is the change in volume (nl), and Δt is the duration of the experiment (min).

## Results

### Leptophragmata require the secondary cell transcription factor Tiptop

To confirm our previous finding that the transcription factor Tiptop is expressed in the leptophragmata of *Tribolium*, in addition to the secondary cells of the free tubule region (17), we made a detailed characterisation of Tiptop expression. Firstly, we assessed the expression pattern of *tiptop* during *Tribolium* embryonic development using fluorescent hybridisation chain reaction (HCR).

We found *tiptop* is first expressed in all cells of the MpTs as they bud from the developing hindgut (Fig. 1A), a result consistent with a previous report (19). Subsequently, as the MpTs begin to grow and extend, we observed that *tiptop* expression refines to their distal region (Fig. 1B). From *Drosophila* we know that the early MpTs which bud out from the developing gut are comprised solely of principal cells (references within 28). Later in embryogenesis, as the MpTs extend, the secondary cells insert into the central MpT region, subsequently dispersing among the principal cells (29, 30). Assuming that the secondary cells in *Tribolium* incorporate into the MpT with a similar timing and distribution, then the *tiptop* expression in the *Tribolium* tubule observed in early and mid-stage embryos is likely to be in principal cells. In late stages of *Tribolium* embryogenesis *tiptop* expression was no longer seen in the distal MpT, but was observed in a subset of cells regularly scattered along the entire MpT length. This includes both the regions associated with the CNC, which we refer to as perirectal tubules, as well as the free tubule regions (Fig. 1C). This could be explained by *tiptop* expression being activated in the secondary cells, and downregulated in the distal principal cells. These distinct phases of *tiptop* expression echo those found for Tiptop in *Drosophila* tubules during embryogenesis (18).

**Figure 1.**
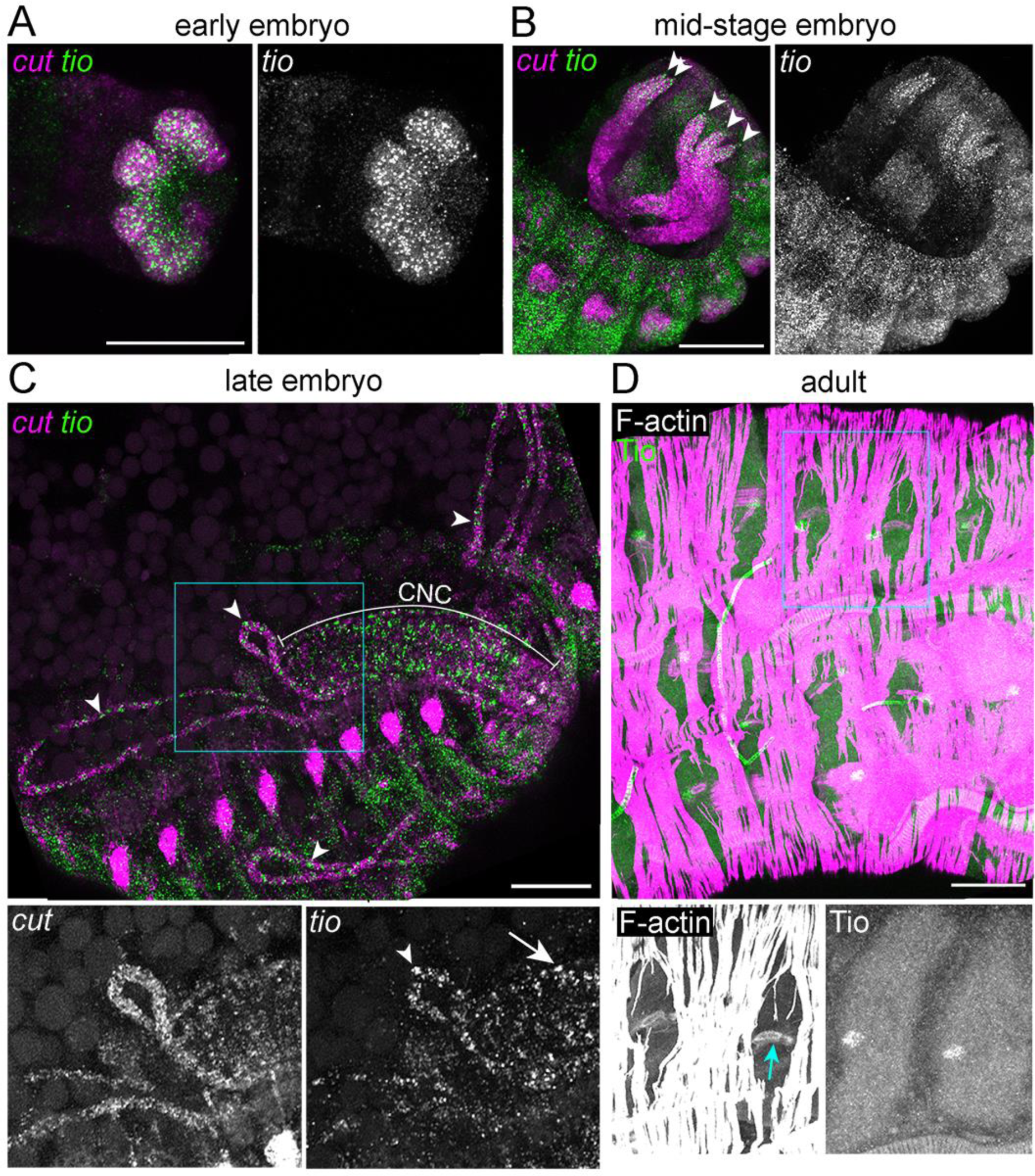
*tiptop* is expressed in leptophragmata. **(A-C)** *Tribolium* embryos stained for *cut* and *tiptop* (*tio*) mRNA. **(A)** An early embryonic stage, in which the six Malpighian tubules (MpTs) have begun budding out from the developing gut. *tiptop* is expressed throughout the MpTs **(B)**. A mid embryonic stage, in which *tiptop* is expressed in the distal segment of the growing MpTs. Arrowheads indicate distal ends of MpTs. **(C)** A late embryonic stage in which *tiptop* is expressed in cells scattered along the free MpTs (arrowheads indicate examples) and perirectal MpTs (arrow indicates example). CNC indicated. Single channel images correspond to region of blue box in merged image. **(D)** Dissected adult CNC stained with phalloidin to reveal F-actin rich structures, and anti-Tiptop (Tio). Blue arrow indicates bar like F-actin structure within leptophragmata. Scale bars in A-C = 50μm and bar in D = 20μm.

Based on the expression pattern of *tiptop* in the perinephric tubules, and on our prior observations (18), we considered these *tiptop* positive cells are likely to be leptophragmata (Fig. 1C). To test this, we dissected adult CNCs and used an antibody to detect Tiptop protein, in combination with counter-staining for F-actin, allowing us to identify the leptophragmata by position and cellular morphology. In *Tribolium* and the related Tenebrionid beetle, *Tenebrio molitor*, the perinephric membrane ensheaths the CNC but is punctuated with small windows under which the leptophragmata sit (5, 6, 14, 16).The outer perinephric membrane, which stains strongly for F-actin, and the windows within it, can readily be identified. The leptophragmata are also apparent, both by their position beneath the window and by the distinctive bar-shaped F-actin structure contained within them. Using this approach, we identified leptophragmata and confirmed that they express Tiptop (Fig. 1D). Together these data show that Tiptop expression initiates in *Tribolium* tubules from embryonic stages and is subsequently maintained in tubule cells (including leptophragmata) into adults, and point towards a likely role in tubule cell identity.

To test for a functional requirement for Tiptop in the development of the leptophragmata, we sought to determine the consequence of *tiptop* depletion. To do this we injected dsRNA targeting *tiptop* into adult females, and observed their offspring at the stage of 1^st^ instar larvae. In control 1^st^ instar larvae we identified leptophragmata based on their expression of Tiptop, their small nuclear size (relative to surrounding principal cells), their position under the perinephric membrane windows and by their F-actin bar structure which, despite being smaller than in the adult, are still discernible (Fig. 2A). Following knockdown, Tiptop expression is abolished (indicating efficient knock-down) and there is no evidence of leptophragmata based on their distinctive morphologies of the F-actin bar structure and small nuclei. (Fig. 2A-B). This suggests that *tiptop* is required for the specification and/or normal differentiation of leptophragmata in the embryo. This is in line with the role for *teashirt*/*tiptop* genes in the differentiation of secondary cells in *Drosophila* (18).

**Figure 2.**
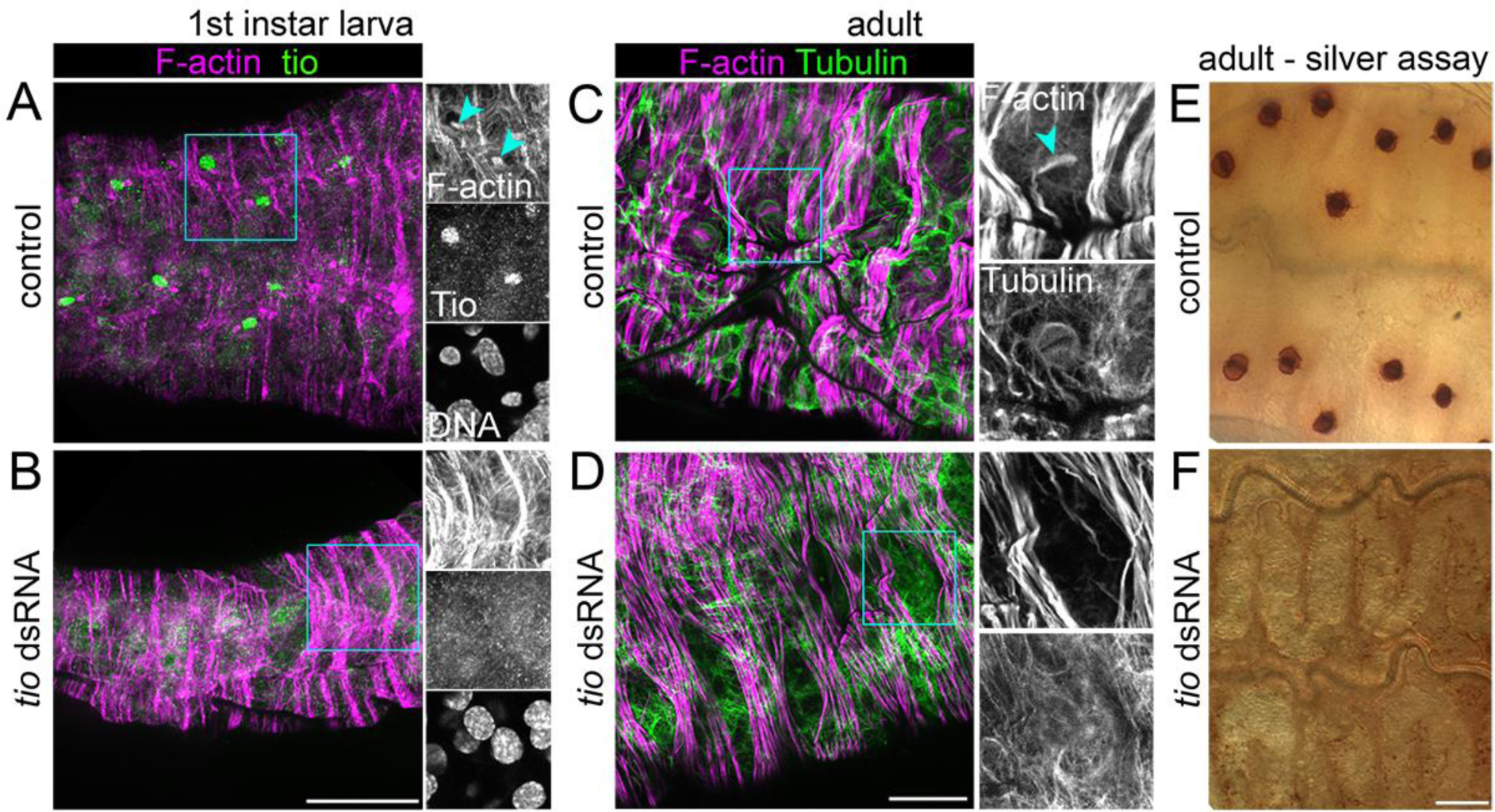
Tiptop is required for leptophragmata development and maintenance. **(A-B)** Dissected 1^st^ instar larval *Tribolium* CNCs stained with phalloidin to show F-actin structures, anti-Tiptop (Tio) and DAPI to label DNA/nuclei. **(A)** Uninjected control. Blue arrowheads indicate actin-rich bars in leptophragmata. **(B)** *tiptop* depleted progeny from adult female injected with dsRNA. **(C-D)** Dissected adult CNCs stained with phalloidin to show F-actin structures and anti-alpha-Tubulin to show microtubule structures. **(C)** Control adult, injected with buffer as a final instar larva. Blue arrowhead indicates actin-rich bar in leptophragmata. **(D)** *tiptop* depleted adult, injected with dsRNA as a final instar larva. **(E-F)** Silver staining of adult CNCs. Precipitated silver (dark regions) is generated at the leptophragmata, indicating abundant chloride ions. **(E)** Uninjected control. **(F)** *tiptop* depleted adult, injected with dsRNA as a final instar larva. Single channel images correspond to region of blue boxes in merged images. Scale bars = 20μm.

To test whether Tio is required for the maintenance of leptophragmata identity we performed *tiptop* knock-down by injecting last instar larvae and analysed adult phenotypes after eclosion. We observed severe morphological abnormalities including deformations in elytra shape and, in the large majority of adults that emerged, incomplete shedding of the pupal case (Fig. S1A-B). The abnormal adults appeared to desiccate rapidly and generally died within a few days of eclosion.

We dissected CNCs from these *tiptop* depleted adults, and found that leptophragmata are lost based on morphological criteria including the absence of the F-actin bar structure and their distinctive staining for microtubules at this stage (Fig. 2C,D). Additionally, we tested for the presence of leptophragmata using an independent physiological assay. Evidence that leptophragmata act as a major site of chloride ion transport was first obtained by incubation with silver nitrate (15). This precipitates as silver chloride in the presence of chloride ions, developing into a dark staining of silver in the presence of light. The darkly stained leptophragmata are completely absent following knockdown of *tiptop* (Fig. 2E-F). As part of a parallel study we also found that staining of NHA1, which we demonstrated to be a cation/proton antiporter expressed in the leptophragmata, is not observed following *tiptop* knockdown (16). Together, these findings suggest that *tiptop* is required to specify and maintain leptophragmata identity including for their normal structure and physiological functions.

**Figure S1.**
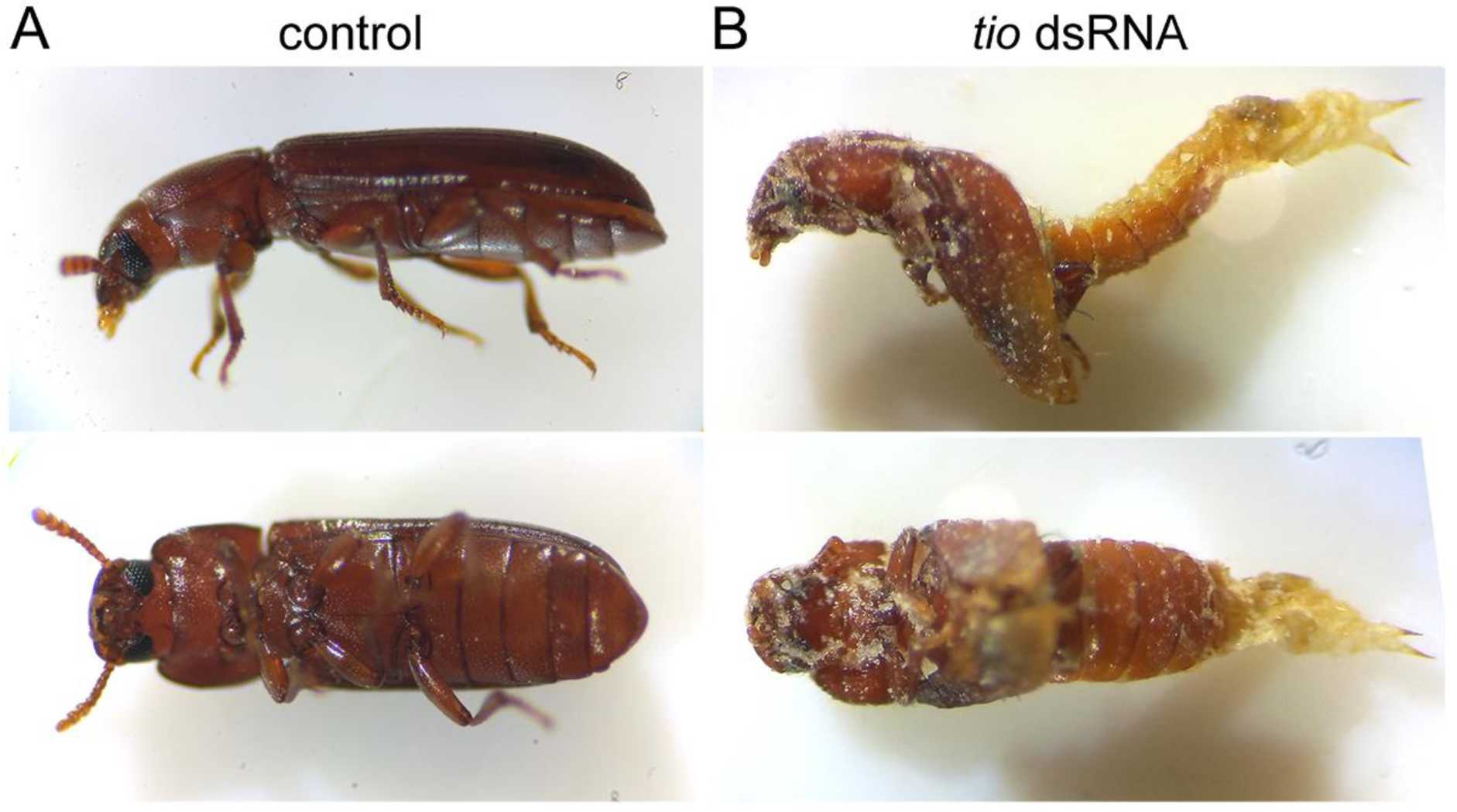
Morphological abnormalities occur during metamorphosis following *tiptop* knockdown. **(A)** Uninjected control adult *Tribolium*. **(B)** *tiptop* depleted adult *Tribolium*, which has been injected with dsRNA as a final instar larva. Upper panels show lateral, and lower panels show ventral view.

### Testing for additional transcription factors required for leptophragmata identity

As both leptophragmata and free tubule secondary cells express Tiptop, we hypothesised that additional transcription factor(s) are involved in regulating leptophragmata versus free tubule secondary cell identity, and further hypothesised these additional transcription factor(s) are also likely to continue to be expressed in the mature leptophragmata, playing a maintenance function as seen for *tiptop*. We therefore decided to carry out a targeted screen to identify such factors, using expression data to guide the selection of candidates.

We first identified >600 transcription factor genes in genome of *Tribolium* (Table S1). These are either orthologues of *Drosophila melanogaster* genes annotated as transcription factors, or genes from the Regulator database of metazoan transcription factors (31; http://www.bioinformatics.org/regulator/). From this list, we then identified genes for which expression was enriched in the CNC relative to other tissues (Table S1). To do so we used BeetleAtlas: a comprehensive bulk RNA-Seq expression database from dissected adult and larval tissues (20, 32)(BeetleAtlas.org). We selected the genes that are most highly enriched in the CNC relative to other tissues, and excluded those where expression is also high in the non-CNC portion of the hindgut, or the free section of the tubule. This allowed us to identify 10 transcription factors with expression most specific to the CNC (Fig. 3).

**Figure 3.**
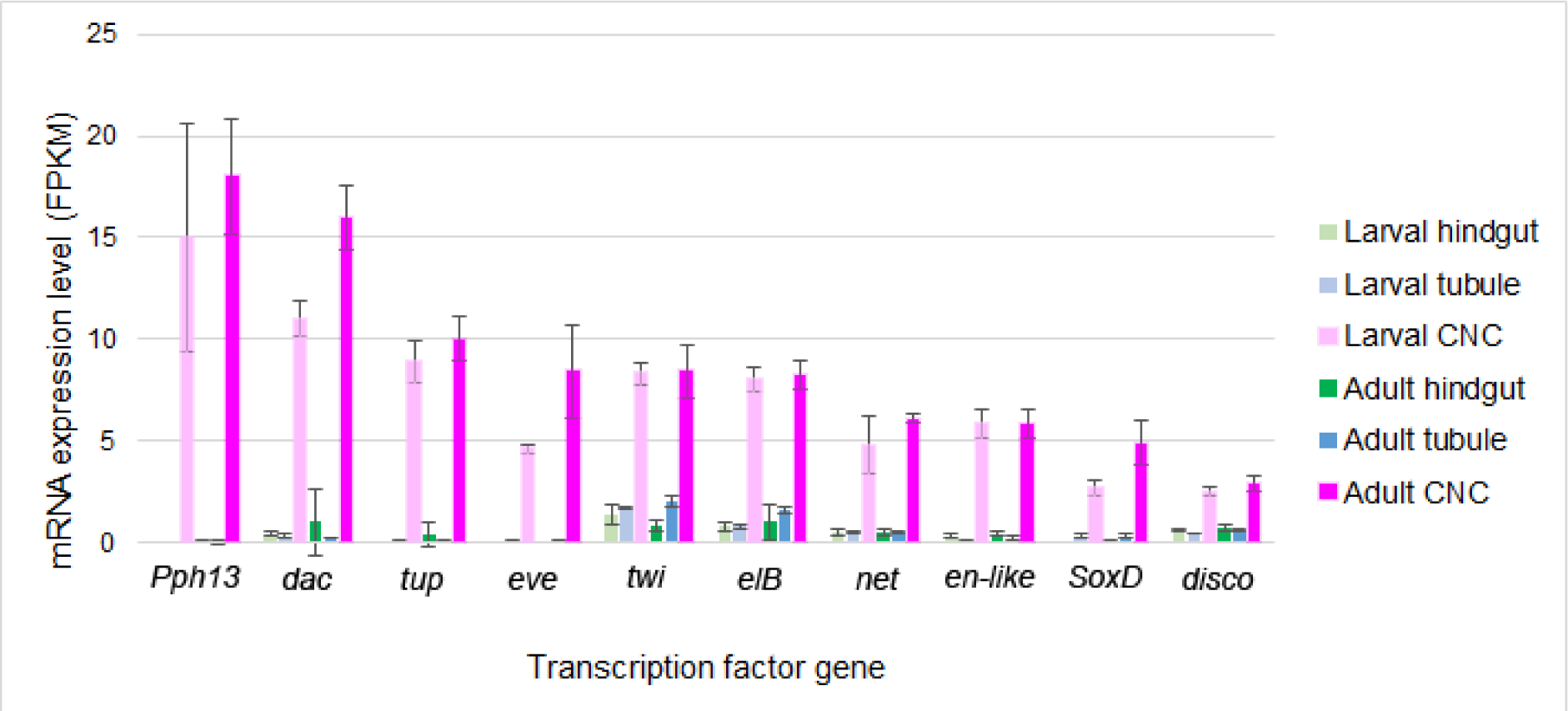
Transcription factors enriched in the cryptonephridial complex. RNA-Seq data from dissected larval and adult *Tribolium* tissues, from BeetleAtlas, showing the mRNA levels of the transcription factors most enriched in the CNC, and comparative levels from the non-cryptonephridial region of hindgut, and from the free region of the MpTs. *Pph13* (*PvuII-PstI homology 13*, TC006110), *dac* (*dachshund*, TC032755), *tup* (*tailup*, TC033536), *eve* (*even skipped*, TC009469), *twi* (*twist*, TC014598), *elB* (*elbow B*, TC000868), *net* (TC005579), *en-like* (*engrailed-like*, TC009896), *SoxD* (TC007419), *disco* (*disconnected*, TC001693).

We injected dsRNA for each gene into final instar larvae and analysed adult CNCs from these animals morphologically, as well as physiologically using the silver staining assay. From this we identified several phenotypes following gene knockdown (Fig. S2, S3 and Table S2).

**Figure S2.**
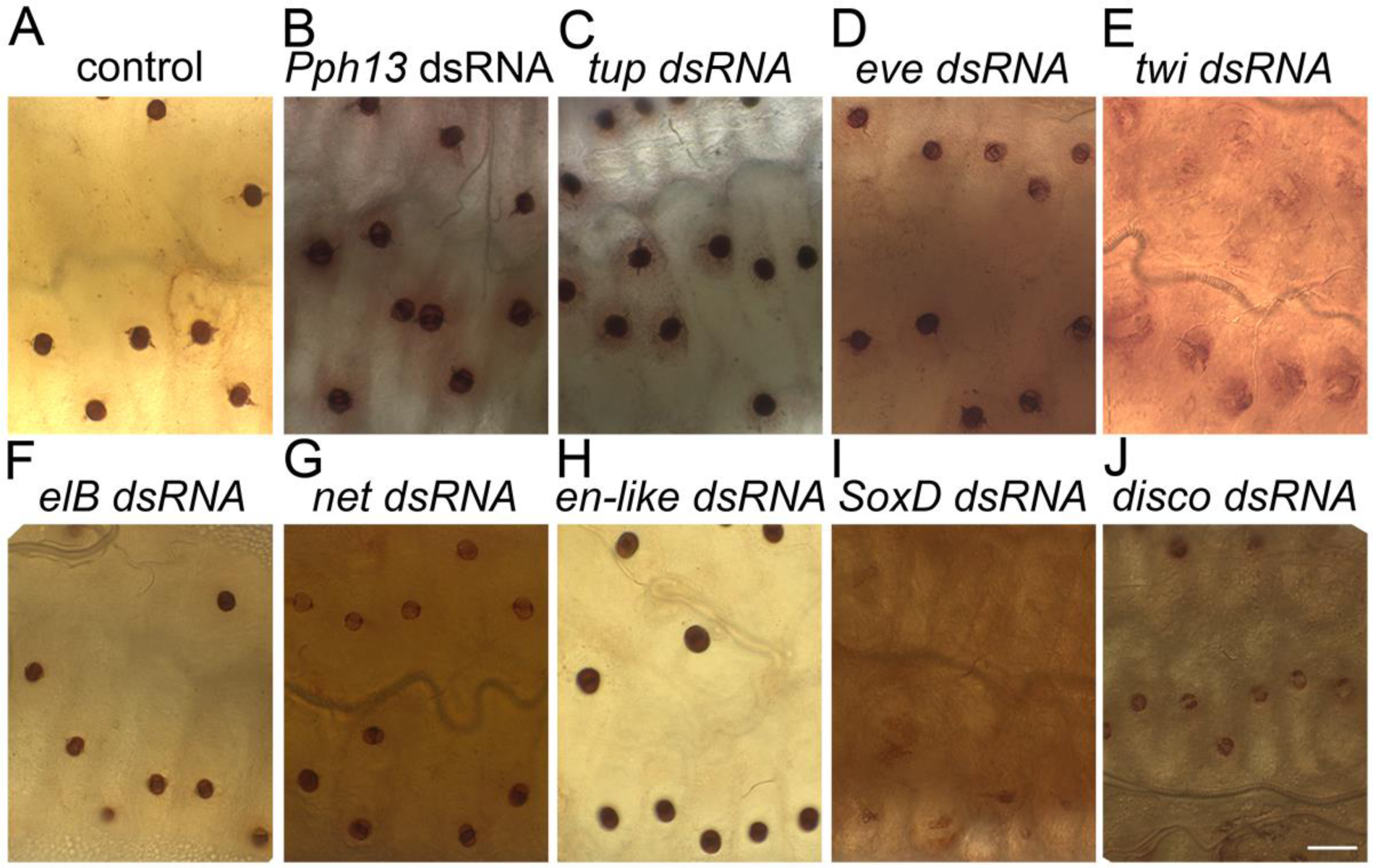
Silver staining assay following knockdown of candidate transcription factors. Dissected adult CNCs incubated with silver nitrate to assay for the presence of chloride ions. **(A)** Uninjected control. **(B-J)** Injected as final instar larvae with dsRNA targeting genes as specified. **(B)** *Pph13*. **(C)** *tup*. **(D)** *eve*. **(E)** *twi*. Leptophragmata are extremely weakly stained. **(F)** *elB*. **(G)** *net*. **(H)** *en-like*. **(I)** *SoxD*. Leptophragmata are extremely weakly stained. **(J)** *disco*. Central region of leptophragmata lack staining. Scale bar = 20μm.

Knockdown of *twist*, *SoxD* and *disco* resulted in weak silver staining in leptophragmata (Fig. S2A,E,I,J and Table S2), suggesting an impaired ability to transport or retain chloride ions. Depletion of *twist* and *SoxD* also resulted in an abnormal structure of the inner layer of the perinephric membrane, which is rich in microtubules. Anti-tubulin staining reveals a region less dense in microtubules surrounding each leptophragmata (Fig. S3A), but this feature appeared to be lost in *twist* and *SoxD* knockdown animals (Fig. S3E and Table S2). FISH-HCR shows these three transcription factors (*twist*, *SoxD* and *disco*) are expressed in the perinephric membrane but not leptophragmata (data not shown), and therefore their effect on the leptophragmata is likely to be indirect. We also found that the leptophragmata appear less well aligned with the windows in the outer layer of the perinephric membrane upon knockdown of *Pph13* (Fig. S3A,B and Table S2), and that the outer layer of the perinephric membrane had a more fragmented appearance following knockdown of *eve* (Fig. S3A,D and Table S2). Although these factors do not appear to be involved in specifying leptophragmata identity, they are candidates for further studies of CNC development and function. For *dachshund* (a transcription factor with well-known developmental roles including in retinal specification and limb patterning (33, 34)), we identified phenotypes which appear to directly relate to a role in the leptophragmata (see following sections).

**Figure S3.**
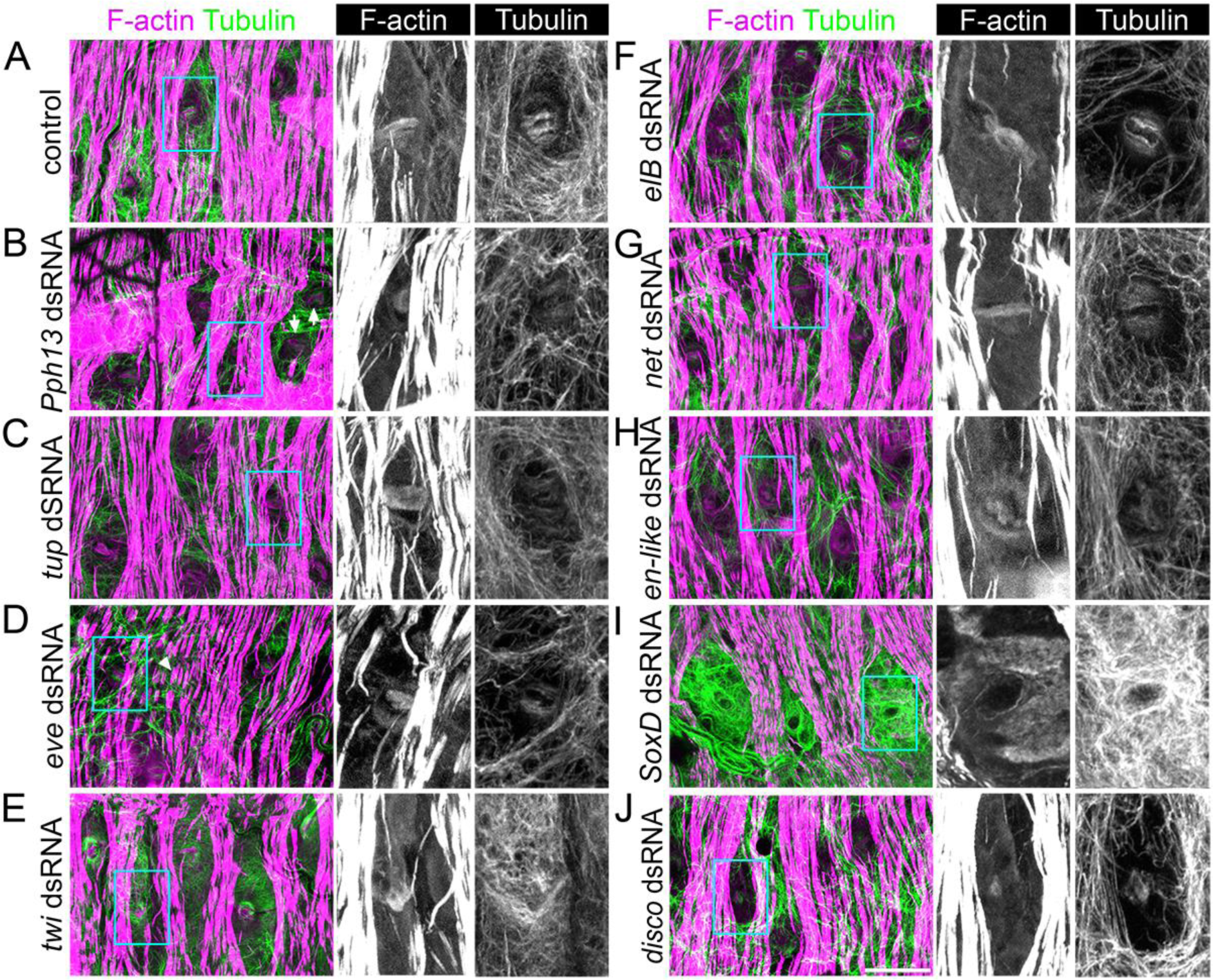
CNC morphology following knockdown of candidate transcription factors. Dissected adult CNCs stained with phalloidin to show F-actin structures and anti-alpha-Tubulin to show microtubule structures. Individuals were injected as final instar larvae targeting genes as specified. **(A)** *GFP* (control). **(B)** *Pph13*. Enlarged panel shows leptophragmata which is largely covered by the outer (actin stained) perinephric membrane. Arrows show two leptophragmata within same window in outer perinephric membrane. **(C)** *tup*. **(D)** *eve*. Arrowhead shows fragmented section of outer perinephric membrane. **(E)** *twi*. No space is apparent between the microtubule rich inner perinephric membrane and the leptophragmata. **(F)** *elB*. **(G)** *net*. **(H)** *en-like*. **(I)** *SoxD.* **(J)** *disco*. Enlarged panel contains example of small/misshapen leptophragmata. Single channel images correspond to region of blue boxes in merged images. Scale bar = 20μm.

**Table S2.** Summary of transcription factor screen phenotypes. Number of CNCs analysed are indicated

| condition | silver nitrate assay phenotype | immunofluorescence phenotype |
| --- | --- | --- |
| control (uninjected) | normal (21), nearly no stain (1) | normal (4) |
| control (*GFP* dsRNA) | normal (6) | normal (5), less dense microtubule region of inner perinephric membrane, which usually surrounds leptophragmata, absent (1) |
| *Pph13* | normal (6), nearly no stain (1) | normal (2), leptophragmata not always aligned with windows in perinephric membrane (4) |
| *tup* | normal (5), leptophragmata appear as empty rings (1) | normal (6) |
| *eve* | normal (5) | normal (1), F-actin staining in outer perinephric membrane separated into blocks rather than continuous bands (4) |
| *twi* | extremely faint staining (5) | less dense microtubule region of inner perinephric membrane, which usually surrounds leptophragmata, absent (8) |
| *eIB* | normal (4), weaker staining (1) | normal (6) |
| *net* | normal (5) | normal (4), less dense microtubule region of inner perinephric membrane, which usually surrounds leptophragmata, absent (1) |
| *en-like* | normal (6) | normal (3), somewhat fragmented appearance of F-actin-stained outer membrane (3) |
| *SoxD* | extremely weak staining (4), somewhat misshapen leptophragmata (2) | less dense microtubule region of inner perinephric membrane, which usually surrounds leptophragmata, absent in all cases, and: F-actin bar in leptophragmata weak and unusually shaped or absent, and outer perinephric membrane fragmented (2), F-actin bar in leptophragmata weak and unusually shaped or absent (3), otherwise normal (1) |
| *disco* | normal (3), faint staining (1), central band of leptophragmata lacks staining (1) | leptophragmata somewhat misshapen, and: perinephric membrane normal (4), inner perinephric membrane abnormal (3), outer perinephric membrane abnormal (2), inner and outer perinephric membrane abnormal (1) |
| *dac* | strongly stained bar across leptophragmata absent (7) | leptophragmata have circular structure enriched in microtubules, F-actin bar in leptophragmata weak and unusually shaped. Nuclei large (9) |

### Identification of Dachshund’s requirement for leptophragmata identity

Leptophragmata were still clearly discernible upon knockdown of *dachshund*, which is in contrast to what we found for *tiptop* knockdown, however their cytoskeletal morphology is dramatically altered: the F-actin bar structure is severely reduced and the microtubules form a knot within the cell which they do not in mock injected animals (Fig. 4A-B and Table S2). Again, unlike *tiptop* knockdown, sliver staining revealed that leptophragmata are present and are physiologically functional, at least in their capacity to transport/accumulate chloride ions. However the darker bar like staining across the leptophragmata, which can be seen in the controls and mock injected animals, was absent (Fig. 4C-D), consistent with the altered cell morphology observed when staining for F-actin.

**Figure 4.**
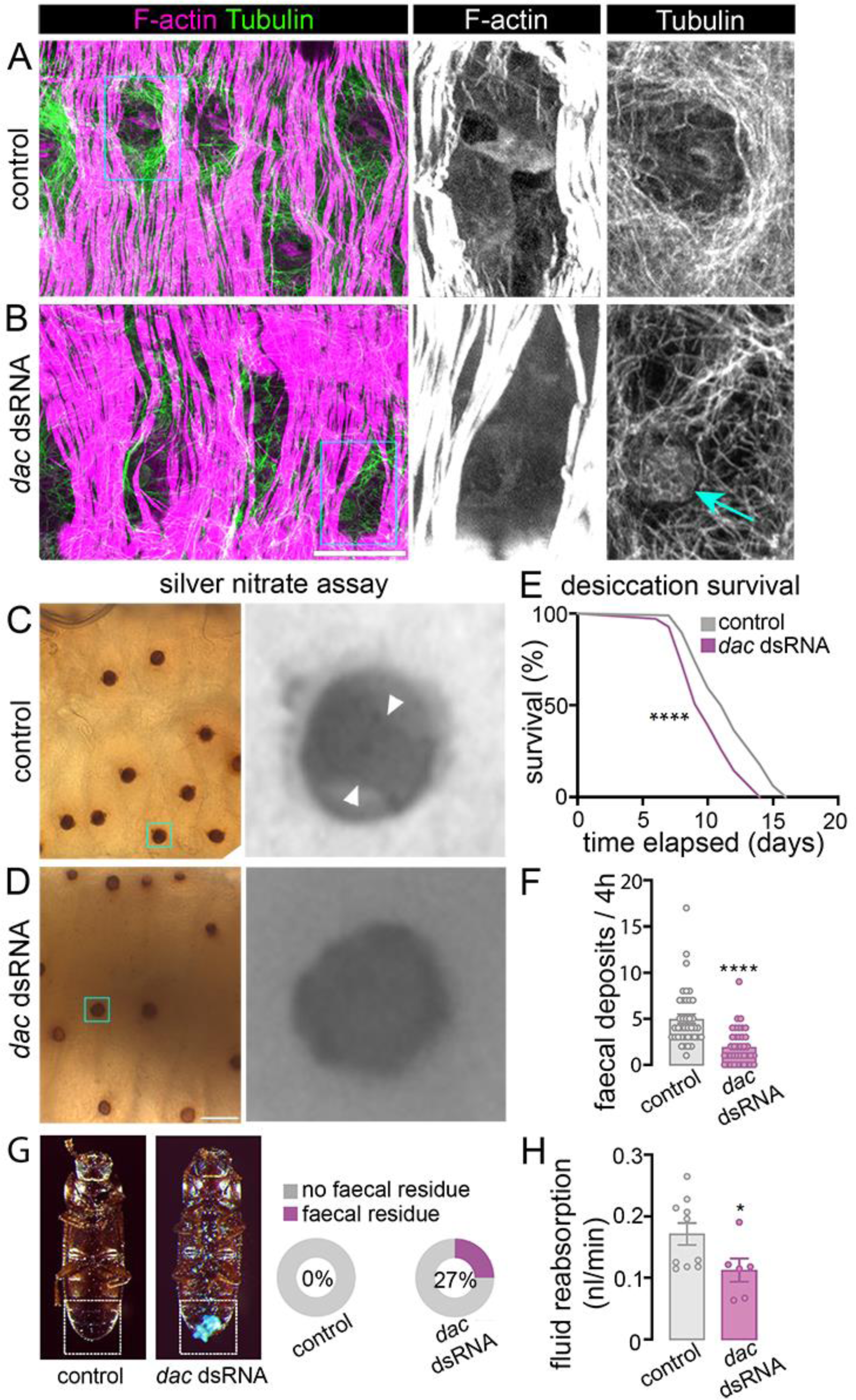
*dachshund* is required for leptophragmata morphology, whole animal water regulation and CNC function. **(A-B)** Dissected adult *Tribolium* CNCs stained with phalloidin to show F-actin structures and anti-alpha-Tubulin to show microtubule structures. Individuals were injected as final instar larvae targeting the genes as specified. **(A)** *GFP* (control). **(B)** *dachshund* (*dac*). Knot of tangled MTs in leptophragmata indicated (blue arrow). **(C-D)** Sliver staining assay in dissected adult CNCs to assay for the presence of chloride ions. **(C)** Uninjected control. Darker stained bar indicated between white arrowheads. **(D)** Injected with *dachshund* dsRNA during the final larval instar. Single channel images correspond to region of blue boxes in merged images. Scale bars = 20μm. **(E)** Kaplan-Meyer survival curves of adult injected with dsRNA targeting *Amp* (control) and *dac* knockdown animals. Knockdown of *dac* reduces organismal survival when exposed to low humidity conditions compared to control (RH 5%, *n* = 69-96, \*\*\*\**P*<0.0001 using log-rank test). **(F)** Defecation rate of *dac* depleted animals is significantly reduced relative to *Amp* dsRNA controls (*n* = 39, **** *P*<0.0001 using t-test). **(G)** Percentage of animals showing faecal residue blocking their rectum for *Amp* dsRNA (control. n=39, and *dac dsRNA* n=48). **(H)** *Ex vivo* preparations of *dac* knockdown beetles show a significantly lower rectal complex-mediated fluid reabsorption rate relative to *Amp* dsRNA control (*n* = 6-10, * *P*<0.05 using t-test).

The majority of *dachshund* knockdown animals survived into adulthood, allowing us to assess both whole organism physiology and, more specifically, CNC function upon Dachshund depletion. We found clear indications that osmoregulation is abnormal. To assess the ability of the CNC to cope under conditions of water scarcity we raised *dachshund* knockdown and mock injected controls under low humidity conditions (∼5% R.H.). We saw a small but significant increase in mortality in the *dachshund* knockdown group (Fig. 4E). In non-desiccating conditions we also identified a reduction in the amount of defecation over a 4-hour period (Fig. 4F). Intriguingly, we also found an increased incidence of animals where faecal residues blocked the anus (Fig. 4G), and speculated this might result from abnormally wet faeces which adhere more readily to the anus, and thus be indicative of increased excretory water loss. Together, these findings suggest altered osmoregulatory performance in the absence of Dachshund, and we decided to assess whether this was a result of perturbed CNC function. To do so we performed an *ex vivo* fluid reabsorption assay using isolated guts cultured under paraffin oil, that we established recently (16). We found a significantly reduction in the rate of fluid reabsorption in preparations depleted for *dachshund* (Fig. 4H). Taken together, these data show that Dachshund is required for normal CNC function and whole animal water regulation.

### Dachshund defines leptophragmata identity

We previously found that the Dachshund protein is expressed in the distal section of the developing MpTs of *Drosophila* (35). Furthermore it has been reported that *dachshund* mRNA is expressed in the distal sections of the developing MpTs in *Tribolium* (36). We used *in situ* HCR to assess *dachshund* mRNA expression in greater detail. Consistent with the previous report, this revealed *dachshund* expression in the distal sections of the developing MpTs (Fig. 5A). In late embryos, *dachshund* expression was restricted to the distal region of the MpT corresponding to the perirectal tubules but was absent from the free tubule region (Fig. 5B). We also sought to determine whether *dachshund* is expressed in the developing leptophragmata in late embryos. By co-staining with *tiptop* mRNA, we found that, *dachshund* is expressed in both principal (i.e. those expressing *cut* but not *tiptop*) and secondary cells (i.e. those expressing *cut* and *tiptop*) in the developing CNC (Fig. 5B).

**Figure 5.**
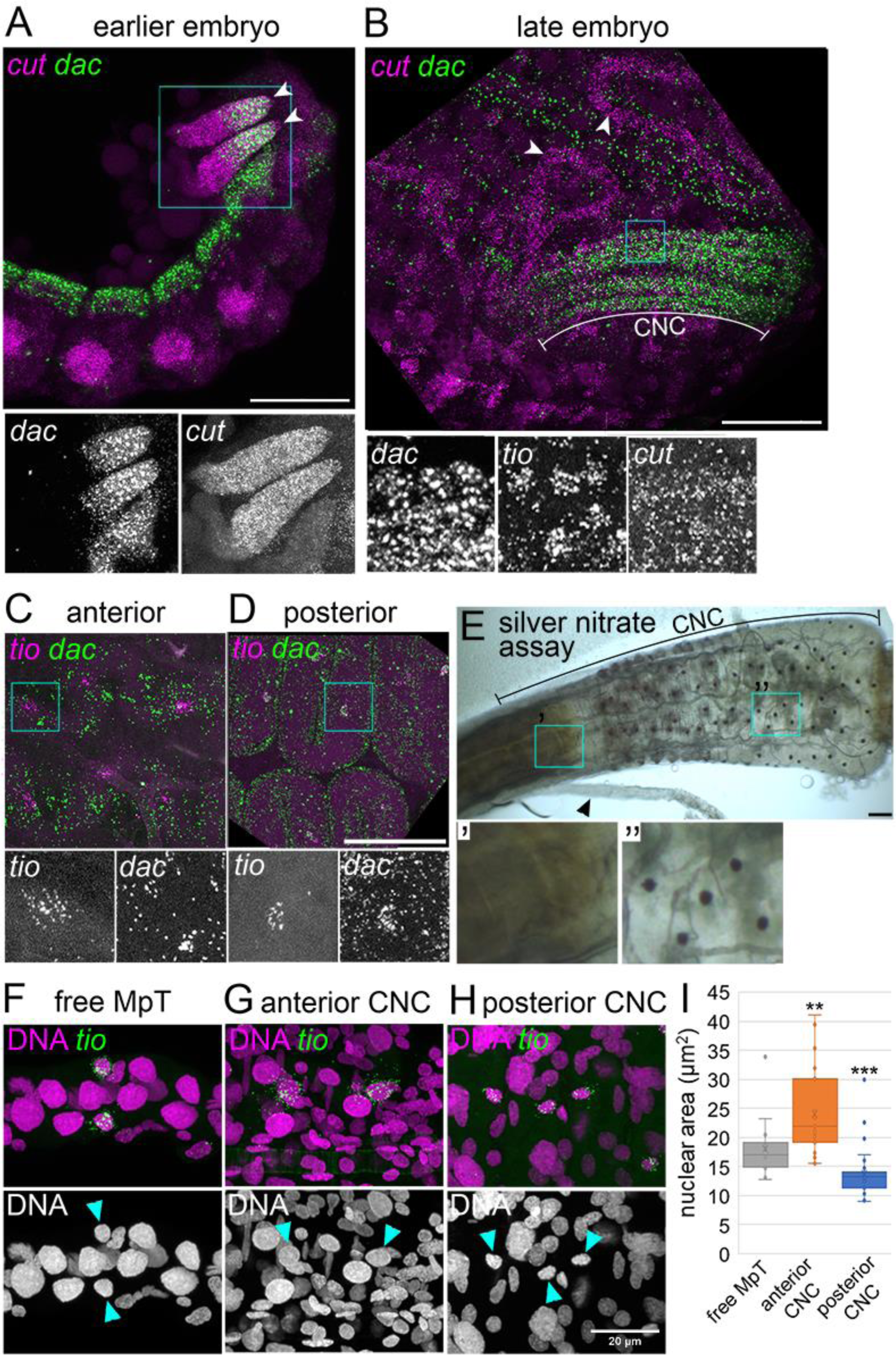
*dachshund* is expressed in the perirectal MpT, including leptophragmata. **(A-D)** HCR-FISH staining of mRNAs as specified. **(A)** Early-stage *Tribolium* embryo stained using probes against *cut* and *dachshund*. Distal ends of MpTs indicated (arrowheads). **(B)** Late-stage embryo stained using probes against *cut*, *dachshund*, and *tiptop*. Free regions of MpTs (arrowheads) and CNC are indicated. **(C-D)** Dissected final instar larval CNCs stained with probes against *tiptop* and *dachshund*. Similar results were observed in adult CNCs (data not shown). **(C)** From the anterior segment of the CNC. **(D)** From the posterior segment of the CNC. **(E)** Silver staining assay in a dissected adult CNC to assay for the presence of chloride ions. Panels below show **(’)** a region of the anterior CNC and **(’’)** a region of the posterior CNC. Enlarged panels below correspond to region of blue boxes in each image. Common trunk where bundle of MpTs enter CNC is indicated (arrowhead). **(F-H)** Nuclei (DNA stained with DAPI) identified for secondary cells (*tio* expressing, determined by HCR-FISH) of the free MpT **(F)**, the anterior CNC **(G)** and the posterior CNC **(H)**. Blue arrowheads identify secondary cell nuclei. **(I)** Adult secondary cell nuclear area for secondary cells of the free portion of the MpT (n=14/9), the anterior CNC (n=23/8, P=0.00228** when compared to free MpT using Mann-Whitney U test), and the posterior CNC (i.e. leptophragmata) (n=32/9, P=0.0006*** when compared to free MpT using Mann-Whitney U test). Sample size values shown as n=number of nuclei/number of animals. Scale bars in A,B and E = 50μm. Scale bar in D and H = 20μm.

**Figure S4.**
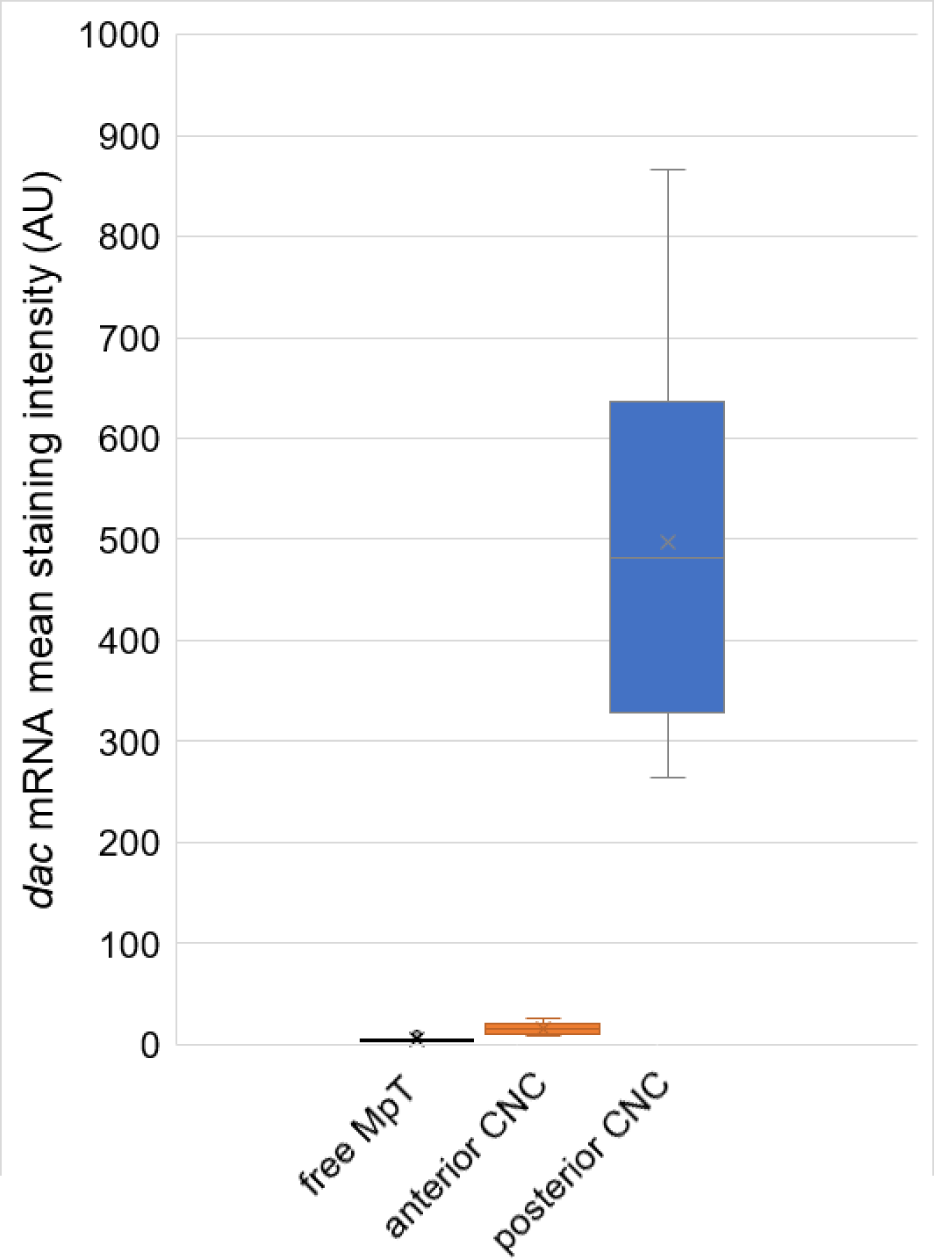
*dachshund* is expressed specifically in the leptophragmata, in contrast to other secondary cells. Fluorescent staining intensity from the HCR-FISH probe against *dachshund* mRNA in secondary cells (including leptophragmata) from regions of the final instar larval free and perinephric MpT, relating to Fig. 5C-D. Free MpT n=8/5, anterior CNC n=12/8, posterior CNC n=13/7. Sample size values shown as n=number of cells/number of animals.

We then defined the pattern of *dachshund* expression in the mature CNC. We found that *dachshund* is expressed in the principal cells throughout the perirectal tubules. We also observed expression of *dachshund* in the *tiptop* expressing leptophragmata within the more posterior part of the CNC. Curiously *dachshund* did not appear to be expressed in the *tiptop* expressing cells within the more anterior part of the CNC (Fig. 5C-D and S4). This prompted us to consider the distribution of leptophragmata in greater detail. We used the silver staining assay, indicating a cell’s ability to transport/accumulate high concentrations of chloride, as a functional marker of the leptophragmata. We used the point where the common trunk of the MpTs joined the CNC as a landmark for the anterior end of the CNC (the common trunk is a segment in which all the free MpTs run together in a single bundle, arrow in Fig. 5E). Using the silver staining assay, we found no stained cells in the anterior portion of the CNC (Fig. 5E’), in contrast to the clearly stained leptophragmata in the more posterior part of the CNC (Fig. 5E’’). We therefore consider that the leptophragmata are exclusively found in the posterior portion of the perirectal tubules, and are distinct from a population of secondary cells in the anterior portion of the perirectal tubules. This is consistent with the lack of leptophragmata previously noted in the anterior CNC region of *Tenebrio* (14). By assessing nuclear size (using DAPI staining) in *tiptop* expressing secondary cells of the perirectal tubules, we also discovered that the nuclei of secondary cells in the posterior CNC are smaller than those of secondary cells in the anterior CNC (Fig. 5F-I), providing an additional marker for leptophragmata and further evidence that leptophragmata are distinct from the secondary cells residing in the anterior perirectal tubules. We observed ∼7-8 anterior secondary cells in each perirectal MpT, and consider these to be a distinct subpopulation of secondary cell which has not been characterised previously, and which appear to differ from the secondary cells of the free MpT in addition to the leptophragmata, as indicated by their different nuclear size (Fig. 5I). The functions of this anterior subpopulation are currently not known.

We wanted to understand further the role Dachshund plays in the specific differentiation of leptophragmata (in contrast to other secondary cell populations), particularly their physiology and endocrine regulation. Our knowledge of specific proteins that define different secondary cell populations in the *Tribolium* renal system, such as differentially expressed ion channels, is limited. However we have shown previously that the hormone receptor Urinate Receptor (Urn8R), which regulates MpT fluid secretion, is specifically expressed in secondary cells within the main segment of the free MpT (4), and could therefore be a potential marker to distinguish secondary cells in different regions.

We assessed the pattern of expression of the Urn8R protein in adult CNCs and free MpTs. We observed expression within secondary cells of the free tubule as reported previously (4; Fig. 6A). We also found that Urn8R is expressed in cells within the anterior part of the perirectal tubules, which correspond to the non-leptophragmata secondary cells identified above (Fig. 6B), but not in leptophragmata (Fig. 6C). On this basis we hypothesised that Dachshund could function to restrict the expression of Urn8R, repressing (directly or indirectly) its expression specifically within the leptophragmata. We therefore looked at expression of Urn8R upon knockdown of *dachshund*. We observed clear expression of Urn8R within leptophragmata, when depleting *dachshund* (Fig. 6D). We reported previously that the diuretic hormone, DH37, signals via Urn8R in secondary cells of the free tubule. Using a fluorescently labelled DH37, we had shown the ability of this hormone to bind free tubule secondary cells (4). We found that the labelled DH37 also binds a population of cells in the anterior CNC (Fig. 6E) likely to be the Urn8R expressing secondary cells, thus indicating that DH37 is competent to bind with the receptor in these cells.

**Figure 6.**
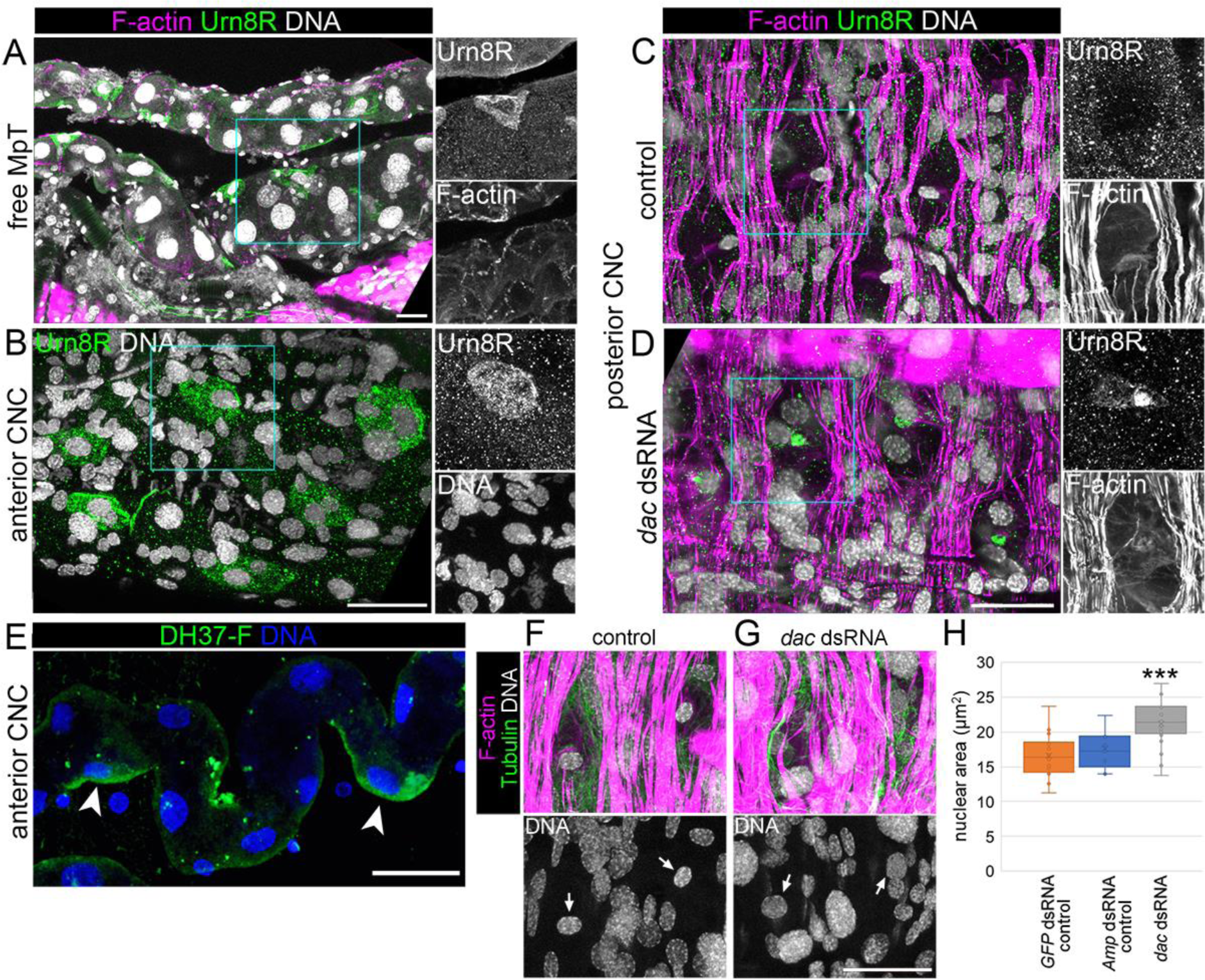
Dachshund is required for leptophragmata differentiation. **(A-D)** Dissected free and perirectal MpTs of adult *Tribolium*, stained with phalloidin to show F-actin structures, DAPI to show DNA/nuclei and anti-Urn8R. **(A)** A region of free MpT from a buffer injected control animal. **(B)** A region of the anterior CNC from a buffer injected control animal. **(C)** A region of the posterior CNC from an uninjected control animal. **(D)** A region of the posterior CNC from an animal injected with dsRNA against *dachshund* as a final instar larva. Single channel images correspond to region of blue boxes in merged images. Scale bars = 20μm. **(E)** Dissected anterior perirectal tubule, incubated with fluorescently labelled DH37 peptide to assess spatial localisation of binding. Scale bar = 30µm. **(F-G)** Dissected adult *Tribolium* CNCs stained with phalloidin to show F-actin structures, anti-alpha-Tubulin to show microtubule structures and DAPI to show DNA/nuclei. Injections of dsRNA against genes as specified was carried out during the final larval instar. **(F)** *GFP* knockdown (control). **(G)** *dachshund* knockdown. Scale bar = 20μm. **(H)** Leptophragmata nucleus area for injection of dsRNA against genes as specified. *GFP* dsRNA (control, n=22/6), *Amp* dsRNA (control, n=9/2), *dac* dsRNA (n=26/8, P=0.00038***). P-value calculated using t-test, comparing to *GFP* dsRNA control, using the mean nuclear area for each animal. Sample size values shown as n=number of nuclei/number of animals.

As a further readout of cell identity, we measured the area of leptophragmata nuclei following *dachshund* depletion, as we had found this parameter to also differ between different secondary cell subpopulations. We found increased area in the *dachshund* knockdown condition. This provides a further indication that Dachshund functions in specification of leptophragmata identity, and loss of *dachshund* possibly results in a shift towards anterior CNC secondary cell identity, which we had found to have a larger nuclear size (Fig. 5G-I).

**Figure S5.**
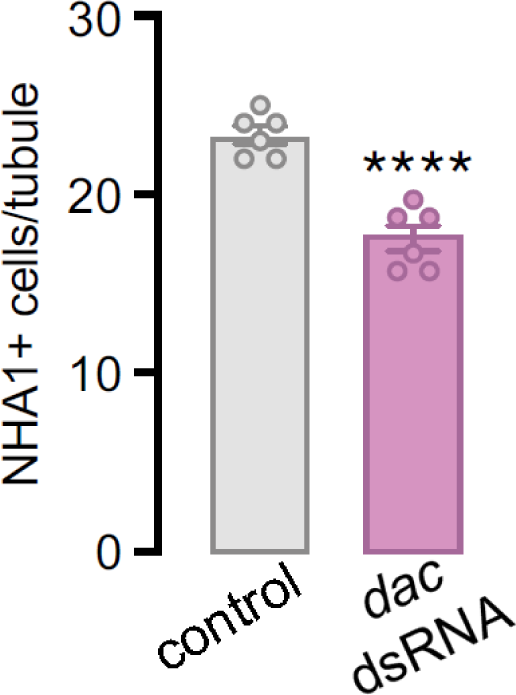
Number of cells staining with anti-NHA1 in the perirectal tubules of *Amp* (control, n=6) and *dachshund* dsRNA injected (*dac* dsRNA, n=6, P<0.0001) perirectal tubules, showing a significant reduction in NHA1+ cells following Dachshund knockdown.

Additionally we assessed expression of the NHA1 protein, which we previously identified as a proton/potassium antiporter, which is expressed predominantly in the leptophragmata (16). By antibody staining we observed NHA1 to still be expressed in leptophragmata following knockdown of Dachshund, although there appears to be a significant change in the number of cells expressing NHA1, with apparent loss of expression in the leptophragmata in the anterior region of the complex (Fig. S5).

Overall our findings suggests Dachshund specifies leptophragmata identity, and in its absence these cells adopt identities more like the other secondary cell populations in the anterior CNC or free tubule.

### Dachshund’s role in distal secondary cell identity is evolutionarily conserved across insects, indicating an ancestral role

The CNC and leptophragmata identity are derived features of the Coleopteran renal system, and it is conceivable that Dachshund was recruited to regulate distal tubule identity during the evolution of the system. However, an argument against this comes from our knowledge that Dachshund is expressed in distal MpT cells in *Drosophila* (an insect without a CNC or leptophragmata) (35). This raises the alternative hypothesis that Dachshund has an ancestral role in patterning distal MpT cell identity across the insects. With this in mind we explored the role of Dachshund in *Drosophila* MpT secondary cell development.

We firstly co-stained *Drosophila* embryos with antibodies against Dachshund, along with Teashirt which is known to mark the secondary cells (18). We found that Dachshund is expressed in the distal, but not the proximal, secondary cells, in addition to the distal principal cells (Fig. 7A). We also looked at the expression of Dachshund in the adult MpTs of *Drosophila*. At this stage, morphologically distinct bar (distal) and stellate (proximal) secondary cells can be identified in the anterior MpT pair. We marked secondary cells (using *tsh-c724-Gal4*), and found that as in the embryo, Dachshund is expressed in both principal and secondary cells. Interestingly, Dachshund is restricted to the bar cell secondary population (Fig. 7B), suggesting it contributes to distinguish bar cell compared to stellate cell subtypes. These findings reveal that Dachshund is expressed similarly in the MpTs of *Tribolium* and *Drosophila*, and suggest that Dachshund could have functioned to specify an ancestral distal secondary cell type which gave rise to the leptophragmata and bar cells in these two lineages.

**Figure 7.**
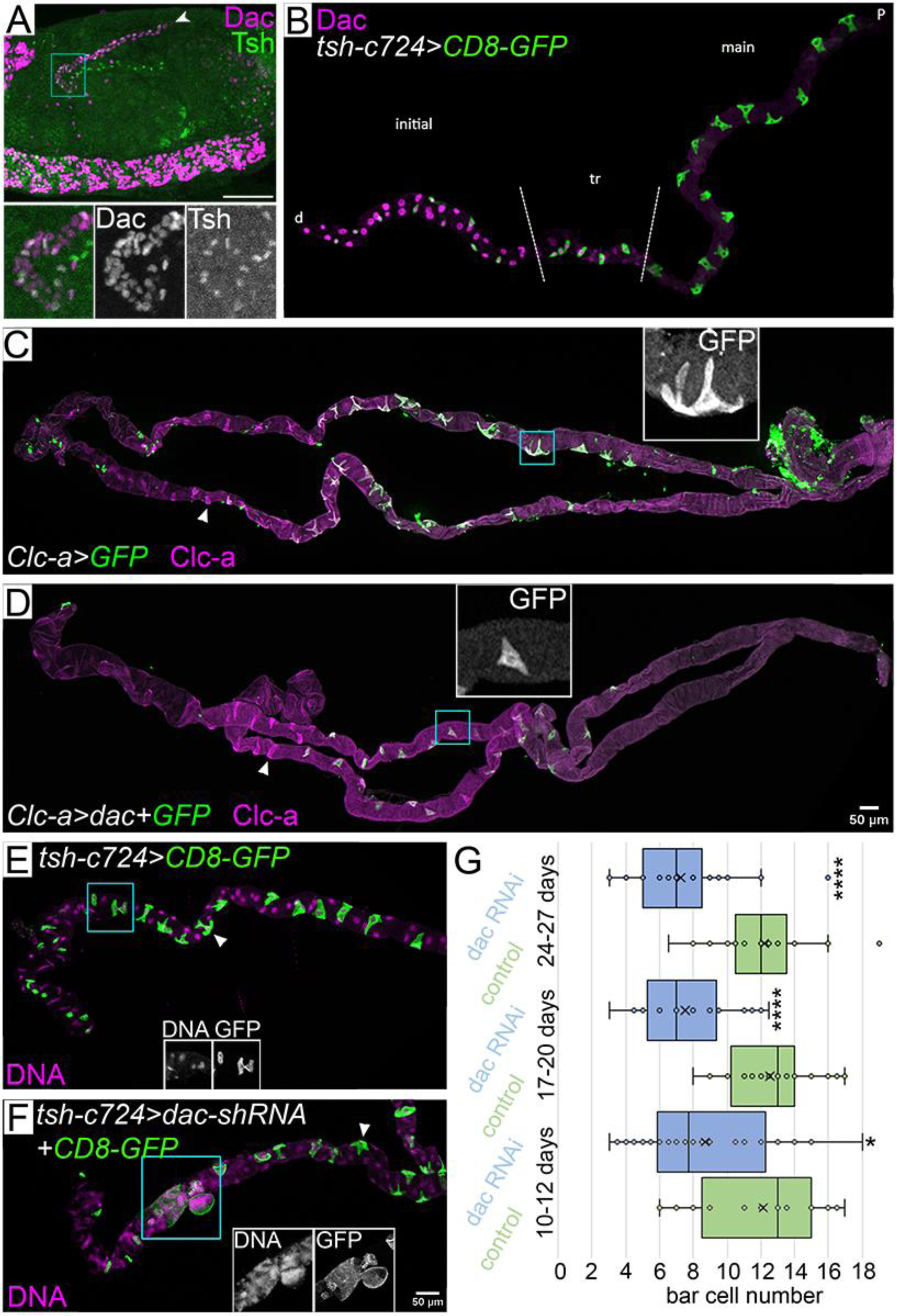
Role of Dachshund in specifying distal secondary cells is conserved in *Drosophila*, which lacks the cryptonephridial arrangement. **(A)** *Drosophila* embryo stained with anti-Dachshund (Dac) and anti-Teashirt (Tsh). Distal end of an anterior MpT marked (arrowhead). **(B)** Dissected anterior MpT from a *Drosophila* adult with GFP-membrane labelled secondary cells (*tsh-c724-Gal4>UAS-CD8-GFP*), stained with anti-Dachshund and anti-GFP. Orientation of MpT indicated (d=distal, p=proximal) and approximate extents of initial, transitional (tr) and main segments marked. **(C)** Anterior pair of MpTs from a control (*Clc-a-Gal4>UAS-GFP(cytosolic)*) *Drosophila* adult, stained with anti-GFP and anti-Clc-a. Inset panel shows enlarged view of stellate cell, corresponding to blue boxed area, and arrowhead indicates a bar cell. **(D)** As in C, but from an adult ectopically expressing *dachshund* in stellate cells (*Clc-a-Gal4>UAS-dac+UAS-GFP(cytosolic)*). **(E)** Distal end of anterior MpT from a control adult (*tsh-c724-Gal4>UAS-CD8-GFP*) matured for 27 days before staining with anti-GFP along with DAPI to show nuclei/DNA. Arrowhead indicates a stellate cell. **(F)** As for E, but from a dac RNAi adult (*tsh-c724-Gal4>UAS-dac-shRNA + UAS-CD8-GFP*). **(G)** Bar cell number from anterior tubules of tsh-c724-Gal4>UAS-CD8-GFP (control) or *tsh-c724-Gal4>UAS-dac-shRNA + UAS-CD8-GFP* (dac RNAi) adults, matured for 10-12, 17-20 or 24-27 days as stated. In each case the n number is for a single tubule, or the mean from an anterior pair of tubules. P values were calculated using Mann-Whitney U test comparing to control at the equivalent time-point, with significance levels adjusted for multiple comparisons using Bonferroni correction. 10-12 days control (n=21), 10-12 days dac RNAi (n=26, P=0.00386*), 17-20 days control (n=28), 17-20 days dac RNAi (n=20, P<0.00001****), 24-27 days control (n=25), 24-27 days dac RNAi (n=29, P=0.00001****). Single channel images correspond to blue boxed regions in merged images. Scale bars = 50μm.

We then aimed to test for functional roles of Dachshund in *Drosophila* secondary cells. *Chloride channel-a-Gal4* (*Clc-a-Gal4*) is expressed in proximal, but not distal secondary cells (i.e. approximately corresponding to the stellate cells). The *Clc-a-Gal4* reporter appears to represent a subset of the endogenous Clc-a expression pattern, as anti-Clc-a stains at least part of the bar cell population, in addition to stellate cells (Fig. 7C). We used *Clc-a-Gal4* to drive expression of Dachshund in the stellate cells, where it would normally be absent. This led to stellate cells losing their normal stellate morphology, becoming smaller and taking on an appearance more reminiscent of bar cells (Fig. 7C-D). This phenotype may be explained by inhibition of stellate cell gene expression by Dachshund, for example of *RhoGEF64c*, which is specifically expressed in stellate cells and the knockdown of which results in a loss of stellate morphology, likely due to altered F-actin regulation (37).

We also carried out the converse experiment in which we depleted Dachshund in the bar cells. To do this we knocked down *dachshund* within all tubule secondary cells by expression of a dsRNA targeting *dachshund*, and assessed bar cell morphology in adult MpTs. Whilst bar cells with relatively normal morphology and distribution were still observed upon *dachshund* depletion in some tubules, abnormal bar cells were also frequently observed, including abnormally large cells, abnormally small cells, and cells which appeared to be extruding from the tubule. We consider it likely these are dying cells, as staining for DNA was very diffuse, indicative of disrupted nuclei that characterise apoptotic cells. Stellate cells were unaffected, in line with a role for *Dachshund* specifically in bar cells (Fig. 7E,F).

We also found a significant reduction in bar cell number in *dachshund* shRNA expressing MpTs at all time points tested. We wondered if bar cells were progressively lost in *dachshund* depleted MpTs, i.e. if Dachshund functions in bar cell maintenance. Although we found a slight tendency towards fewer bar cells as adult MpTs matured, the difference between timepoints is not significant (Fig. 7G). Overall our findings indicate that Dachshund confers a distal/bar cell identity to the secondary cells in *Drosophila* MpTs, akin to its role in conferring leptophragmata identity in *Tribolium*. Furthermore, it provides evidence to homologise leptophragmata and bar cells, and supports the idea that *Dachshund* confers distal secondary cell identity across insect MpTs.

## Discussion

### Tiptop and Dachshund establish leptophragmata identity

Leptophragmata are a derived and highly specialised cell type of the CNC, whose activities as the main site for ion exchange drive CNC function (16). Here we show that leptophragmata constitute a subpopulation of MpT secondary cells, a cell type found widely in insect MpTs, including those species without a CNC. We find leptophragmata identity is established by the transcription factor Tiptop, akin to the requirement for Teashirt/Tiptop transcription factors for secondary cell identity in *Drosophila* (18). Leptophragmata are distinct from other secondary cells in *Tribolium*’s free MpT region and those in the anterior perirectal MpT, as well as from secondary cells of other species, both in their morphology and function. This posed the question of how different secondary cell subtypes, including the leptophragmata, the secondary cells in the anterior CNC and free MpT region, are established in *Tribolium*, and how they have evolved between species. A screen for further transcription factors that control leptophragmata differentiation identified a role for Dachshund. We find Dachshund controls features of leptophragmata morphology, such as their cytoskeletal arrangement and nuclear size, and acts to regulate gene expression which distinguishes leptophragmata from other secondary cell subtypes. Loss of Dachshund leads to loss of leptophragmata characteristics and their adoption of characteristics resembling the non-leptophragmata secondary cell subtypes. This includes de-repression of the diuretic hormone receptor Urn8R, which is expressed in secondary cells of the free tubule (4) and anterior CNC, but not leptophragmata.

The upstream developmental mechanisms by which these transcription factors become expressed await further investigation. In *Drosophila*, the unique molecular identity of secondary cells, including their expression of Teashirt and/or Tiptop, is a consequence of their distinct developmental origin compared to that of principal cells. The secondary cells in *Drosophila* originate from a subpopulation of caudal visceral mesoderm; they integrate into the developing MpTs and begin to express Teashirt (29, 30). It seems likely that a common developmental process is conserved in *Tribolium*. There is less evidence of how Dachshund’s expression comes to be patterned in the secondary cells. In *Drosophila*, specification of distal MpT identity (including expression of Dachshund) occurs very early in principal cell precursors under the control of Wingless/Wnt signalling, which is expressed asymmetrically in the principal cell precursors (35). It is conceivable that secondary cell identity (in both *Drosophila* and *Tribolium*) could be determined according to whether secondary cells integrate into a Dachshund positive or negative population of principal cells, possibly through interactions with their new principal cell neighbours. Such an interaction has precedent; apical-basal polarity of the principal cells is required for correct polarisation of the secondary cells in *Drosophila* (30).

Do Tiptop and Dachshund function as part of a single transcriptional complex in defining leptophragmata identity? From *Drosophila* it is known that Teashirt/Tiptop transcription factors regulate different sets of genes in a context dependent manner, depending on the presence of other transcriptional coregulators (38–42). Interestingly Dachshund has been shown to block the transcription of a target gene of the Teashirt-Yorkie-Homothorax complex, in the *Drosophila* eye disc (43, 44). It is conceivable that a similar regulation occurs in MpT secondary cells and leptophragmata: Tiptop may activate *urn8r* transcription (and possibly other targets too) in secondary cells of the free tubule and anterior CNC, with Dachshund acting to repress this specifically in leptophragmata. This type of cross regulation might also extend to the control of genes underpinning the distinct morphology of leptophragmata.

### Dachshund regulates the physiological functions of the CNC

Our data suggest that Dachshund regulates the transcriptional control of *urn8r*, and Dachshund may regulate other genes expressed in leptophragmata that underpin their unique morphology and functions. A central role for Dachshund’s regulation of physiologically important targets in the CNC is supported by the *dachshund* knockdown experiments, which lead to defects in CNC osmoregulation and a failure of the animal to tolerate desiccation stress (Fig. 4E-H). However, it is likely other transcription factors act in parallel to Dachshund, as some features of leptophragmata identity remain upon Dachshund depletion, such as their ability to accumulate/transport high levels of chloride ions (Fig. 4D). We are now working to resolve mRNA expression data of the CNC at single cell resolution, to identify further regulators of leptophragmata identity. Further, whilst a role for Dachshund in leptophragmata is clear, Dachshund is also expressed in perirectal principal cells, indicative of a requirement for Dachshund in this subpopulation of principal cells. It remains to be resolved whether defects in osmoregulation and tolerance to desiccation stress seen in the absence of Dachshund reflect a requirement in leptophragmata and/or principal cells.

Why expression of the Urn8R hormone receptor would be actively repressed within the leptophragmata remains another interesting question for future investigation. Distinct physiological functions of insect tubules, including calcium homeostasis, fluid secretion and reabsorption are segregated to discreet regions along their length (21, 45-47). Decoupling of endocrine control of secondary cell activity within different regions might provide a finer level of control. This could be even more important in species with CNCs such as *Tribolium*, where the perinephric MpTs serve a highly specialised function. Our previous results show that activating the Urn8R pathway in free tubule secondary cells has a diuretic effect by stimulating fluid secretion into the MpT lumen, which is then expelled into the gut (4). It would make sense to prevent simultaneous upregulation of secretion by the perirectal MpTs, as this would recycle fluid from the rectum back to the haemolymph, thereby counteracting the diuretic effect. This may thus explain why it is necessary to repress Urn8R expression in the leptophragmata. Our data do not address whether distinct endocrine signals regulate leptophragmata activity but, given the high metabolic costs of the system, it is likely that activity is regulated in alignment to the physiological needs of the animal. Single cell expression data would also help identify candidate endocrinological control mechanisms.

### Evidence that leptophragmata originated from an ancestral distal secondary cell

There is growing evidence that the physiological functions of beetle MpTs are radically different compared to other insects, of which the emergence of the specialised leptophragmata cell is one aspect. These evolutionary changes are likely to underly the ability of some tenebrionid beetle species to survive in very dry environments, so contributing to their remarkable evolutionary success. It is known that major changes to the endocrinological control of MpT cell types have occurred in beetles, including the loss (at genome level) of the kinin hormone pathway that regulates secondary cells in other species, and the restriction of another hormonal pathway active in all principal cells in other species, to just a small subset of principal cells in beetles (3). In the case of the Urn8-receptor pathway, we recently found that the diuretic hormones DH37 and DH47 signal to the free tubule secondary cells to stimulate secretion in beetle species of Polyphaga, including *Tribolium* (4). However, in the more basal beetle group Adephaga, and in other insects such as *Drosophila*, there are indications this pathway instead controls principal cell activity (4, 48).

These evolutionary changes also appear to relate to the movement of water. It is known for many insects that secondary cells are a major site of water flow, facilitated by expression of aquaporin water channels (2, 49). By contrast, leptophragmata are considered impermeable to water; an important feature to prevent water being drawn from the haemolymph into the CNC, which would counter the CNC’s ability to establish a high ionic concentration (14, 50).The ability to transport water may have been specifically lost in leptophragmata, possibly by Dachshund acquiring the ability to inhibit aquaporin gene transcription. There is some evidence that water transport mechanisms may have been altered or lost altogether from the secondary cells of beetles (2, 51), and future analysis of aquaporin expression in *Tribolium* should help resolve this.

Although cellular function has clearly changed dramatically in beetles, we find evidence that the same fundamental cell types are present within the MpTs of *Tribolium* and *Drosophila* (*Drosophila* MpTs appear to be much closer to the ancestral condition). Furthermore, the differentiation of MpT cell types, and the maintenance of their identity, appear to be controlled by the same core transcriptional regulators in these different species. For example, Cut, Teashirt/Tiptop and Dachshund are expressed in the principal cells, secondary cells and distal MpT respectively, with these factors even undergoing similar changes in their expression patterns during development (17, 18, 35). In both species, Teashirt/Tiptop is expressed in all secondary cells, and Dachshund is expressed in a distal subtype of these cells (leptophragmata in *Tribolium* and bar cells in the tubules of *Drosophila*). This suggests that the leptophragmata originated from a secondary cell subtype already present in the ancestral insect, and that the core transcription factors specifying the identity of this cell type have been conserved. Changes to the molecular networks downstream to these transcription factors, however, are likely to have been significantly remodelled, underpinning the evolution of the unique characteristics of leptophragmata. Whilst *Drosophila* bar cells are genetically distinct from the more proximal (stellate) cells (21), we have little understanding of the ancestral or extant functions of the distal secondary cells for any insect. It is known that the initial segment of the MpT, in which the bar cells reside, functions in calcium homeostasis (45, 52), and the specialised role of the bar cells could relate to this.

Our findings provide insights into the evolutionary processes which have given rise to an unusual renal cell in a major beetle lineage. More broadly, this can inform ongoing debates into how novel cell types arise, and supports the paradigm that cell types can be defined by the core set of transcription factors they express, which tend to be conserved over evolutionary time (53–56). In contrast the downstream pathways which ultimately confer phenotypic differentiation of the cell appear to be highly labile to evolutionary change. This study uncovers the developmental mechanisms underlying the establishment of cell identity for a novel renal cell in the beetle CNC: leptophragmata. Leptophragmata are derived cell types that have evolved novel characteristics, underpinning CNC function. Our finding that leptophragmata are derived from ancestral MpT secondary cells whose identity but not downstream characteristics is established by the same patterning mechanisms found in insects without leptophragmata, sheds light on how novel cell types evolve.

## Acknowledgements

We thank Matt Benton for the *Tribolium* stock, the Bloomington *Drosophila* Stock Center and Resource Center (National Institutes of Health grants P40OD018537 and 2P40OD010949-10A1) and the Vienna *Drosophila* Resource Center (VDRC) for *Drosophila* stocks, Julian Dow and Anthony Dornan for *Clc-a-Gal4* and anti-Clc-a, Anisha Kubasik-Thayil (IMPACT imaging facility, University of Edinburgh) for assistance with imaging. This work has been funded by The Leverhulme Trust [RPG-2019-167], The Carnegie Trust Grant [70425], the Biotechnology and Biological Sciences Research Council [BB/N001281/1] and [BB/X014703/1], the Villum Fonden [15365], by the Danish Council for Independent Research [9064-00009B], the Ragna Rask-Nielsens Foundation [KH0622] and research infrastructure grant by Carlsbergfondet [CF19-0353]. For the purpose of open access, the author has applied a CC BY public copyright licence to any Author Accepted Manuscript version arising from this submission.

